# Hippocampal-prefrontal communication subspaces align with behavioral and network patterns in a spatial memory task

**DOI:** 10.1101/2024.07.08.601617

**Authors:** Ryan A. Young, Justin D. Shin, Ziyi Guo, Shantanu P. Jadhav

## Abstract

Rhythmic network states have been theorized to facilitate communication between brain regions, but how these oscillations influence communication subspaces, i.e, the low-dimensional neural activity patterns that mediate inter-regional communication, and in turn how subspaces impact behavior remains unclear. Using a spatial memory task in rats, we simultaneously recorded ensembles from hippocampal CA1 and the prefrontal cortex (PFC) to address this question. We found that task behaviors best aligned with low-dimensional, shared subspaces between these regions, rather than local activity in either region. Critically, both network oscillations and speed modulated the structure and performance of this communication subspace. Contrary to expectations, theta coherence did not better predict CA1-PFC shared activity, while theta power played a more significant role. To understand the communication space, we visualized shared CA1-PFC communication geometry using manifold techniques and found ring-like structures. We hypothesize that these shared activity manifolds are utilized to mediate the task behavior. These findings suggest that memory-guided behaviors are driven by shared CA1-PFC interactions that are dynamically modulated by oscillatory states, offering a novel perspective on the interplay between rhythms and behaviorally relevant neural communication.

## INTRODUCTION

Cognitive functions, including memory and decision-making, hinge on the coordinated interaction across multiple brain regions. In the hippocampus, such coordination prominently anchors to hippocampal oscillatory rhythms (Colgin 2016; Zielinski, Tang, and Jadhav 2020). These rhythms have been hypothesized to help synchronize downstream target networks by aligning periods of excitability (McKenzie 2018). Parallel to this idea, other communication hypotheses have been put forth to explain tightly regulated neuronal communication across regions, including communication subspaces (Kohn et al. 2020; Semedo et al. 2020) and communication through coherence (Fries 2015). Yet it remains unclear how these seemingly parallel aspects of communication—rhythms and alignment of firing to communication subspaces—could relate to one another.

Hippocampal activity in rodents during behavior and sleep displays a diverse cast of high- and low-frequency events. At the high-frequency end, sharp-wave ripples (SPW-R) constituting some of the most synchronous events in the brain occur during periods of immobility and during non-REM sleep (Buzsáki 2015; Zhang et al. 2021). These transient high-frequency bursts have been implicated in interregional memory consolidation and retrieval (Battaglia et al. 2011). During quiescent periods, SPW-Rs are known to mediate interactions with extra-hippocampal areas, including the prefrontal cortex and subcortical structures (Abadchi et al. 2020; Kaplan et al. 2016; Nitzan et al. 2022; Norman et al. 2021), comprising key times of inter- and intra-areal messaging. In contrast, lower frequencies, in particular theta rhythms, predominate during active exploration, motor behavior (Joshi et al. 2022) and memory processing (Seger et al. 2023; Shimbo, Izawa, and Fujisawa 2021). Theta oscillations and theta phase-modulated neuronal activity have been causally linked to memory encoding and retrieval (Siegle and Wilson 2014). Theta may therefore enable and invoke these separate processes in part through regimented and unique timing of inter-areal inputs and outputs within each theta wave (Bezaire et al. 2016; Fernández-Ruiz et al. 2019; Mysin 2023; Tsodyks et al. 1996), possibly linked to gamma rhythms (Colgin et al. 2009; Fernández-Ruiz et al. 2017). Hippocampal spiking thus is known to interact with oscillation timing and vice versa.

Apart from oscillations, communication fundamentally also arises from structural possibilities available in the network, giving rise to only limited manifolds that do not span all possible firing configurations. Studies using recurrent networks have provided valuable insights into communication manifolds. Sparse connectivity creates low-dimensional dynamics (Herbert and Ostojic 2022), and coupling groups of such recurrent networks through sparser inter-network projections creates even lower dimensional structures, a low-rank manifold where activity passes between networks (Barbosa et al. 2023; Kozachkov, Ennis, and Slotine 2023). These narrow-dimension dynamics confer advantages like enhanced stability and prescribe a limited range of spiking patterns when pairs of networks engage in communication (Herbert and Ostojic 2022).

A critical insight is that linear subspaces can approximate variations on nonlinear manifolds, assuming the space is not overly curved (Gallego et al. 2017; Johnson et al. 2003; Semedo et al. 2019). Using such a linear approximation, Semedo et al. (2019) made important discoveries about visual cortical communication, namely, inter-areal V1-V2 interactions occurred in a lower dimension than intra-V1 communication. This V1-V2 subspace was consistent across stimuli, unaffected by nonlinearity, and distinct from part of V1’s subspace for private V1-to-V1 messaging (Iyer et al. 2021; Semedo et al. 2019). These principles have been substantiated and expanded in other work. For instance, Srinath, Ruff, and Cohen (2021) found cortical communication geometry was largely stable when attention shifted, though information flow increased along it, and MacDowell et al. 2023 later expanded the investigation to subserve an area’s multiple possible downstream targets. With large-scale recording, they discovered subspaces projecting to a region’s targets were often multiplexed and overlapping. Thus, as suggested by models (Barbosa et al. 2023), a population can send to or receive from multiple targets and sources by aligning spiking activity to the appropriate subspace (MacDowell et al. 2023; Semedo et al. 2019).

These two forms of communication, rhythmic coordination and subspace alignment, have been well-described. However, whether and how these two communication modes fit together is not clear. Are they separate mechanisms – or intertwined, different perspectives on a shared process? In 2023, Kim et al. observed one possible interaction. In a motor learning task, they found prefrontal (PFC)-motor cortical (M1) communication subspace aligned activity intensifies when M1 slow oscillations (SO) follow hippocampal sharp-wave ripples. In other words, complexes of hippocampal SPW-R and cortical SO mark time periods where PFC and M1 share activity along their communication space. This exemplifies a possible rhythmic event entangling communication spaces. Theoretically, neural manifolds can exhibit limit cycles and rotations in spiking, which could evolve within or between communication and private subspaces or be completely separate. However, direct insights remain scarce. Lastly, communication subspaces have been characterized in response to perceptual stimuli but remain unexplored for cognitive and task-related behaviors.

Since hippocampal activity patterns exemplify rhythmic communication and with evidence that the task under study depends on hippocampal-prefrontal communication (Maharjan et al. 2018), we decided to address these gaps by investigating the following questions: are hippocampal-prefrontal communication subspaces related to task behavior and how do rhythmic network patterns sculpt them? We also asked if these relationships evolve with learning. Clarifying the linkage between rhythms and subspaces could reveal how neural populations temporally harmonize and multiplex communication during cognitive processing.

## RESULTS

Cognitive processes arise from interconnected brain areas working together. A notable example is memory-guided decision-making, a cognitive process shaped by the synchronized relationship between hippocampal area CA1 and PFC. To probe this coordination, we simultaneously recorded CA1 and PFC activities as rats learned a spatial memory task on a continuous W-maze alternation task (**Figure 1A**). The rats were taught to navigate varying paths on a W-maze across 8 distinct training sessions interleaved with rest sessions within a single day (**Figure 1B**, Shin, Tang, and Jadhav 2023). This single-day learned task structure permitted us to simultaneously monitor the activity of the same CA1 and PFC ensembles throughout the course of learning. For success, this required rats to remember their most recent left/right choice while traversing from the central arm in order to choose the opposite side arm (outbound working memory task), and subsequently, to return to the center arm to receive a reward (inbound). As rats learned and performed the task, we monitored the dynamics of cell firing and local field potentials. This yielded 394 CA1 cells and 216 PFC neurons (**Table 1**) using our selection criteria (Methods) across N=7 animals for all analyses unless otherwise noted.

### Communication-aligned activity drives robust, repeatable behavior responses

Our first aim was to examine whether memory-guided behaviors in the task coordinate with communication-subspace-aligned activity. The fundamental question in communication subspaces is to find the changes in target brain area *Y* that arise with changes in a source area *X*. From the point of view of a linear model, this can be approximated with *Y* = *XB*. The instructive power of this technique lies in two observations. If we set *XB*′ = *Y* = 0 and solve for *B*′, the coefficients encode a special solution where source neuronal activity is insulated from influencing the target area Y (Kaufman et al., 2014); this is called a private space. Conversely, solving instead for *XB* = *Y*, the resulting remaining dimensions of the source neurons approximate the activity space of *X* that change *Y*, hence the shared communication subspace (schematic in **Figure 1 – Supplement 1**). This is illustrated in **Figure 1C** where we schematize a quad-neuron system with 3 CA1 source neurons and 1 PFC target neuron. An example of a 1-dimensional communication subspace is shown via a line where source neuron activities comodulate with a target neuron (Semedo et al. 2019). Directions off from this axis are private: signals in the source area can fire without influencing the target, insulating it from communication between the networks. When many such target neurons are measured in this fashion (**Figure 1D**), it comprises a collective communication subspace. We wished to understand how and if these spaces relate to memory-guided behavior, a hallmark of CA1-PFC network functions, and rhythmic network patterns, believed to be critical in CA1-PFC interaction (Floresco, Seamans, and Phillips 1997; Maharjan et al. 2018; Zielinski et al. 2020).

**Figure 1.**
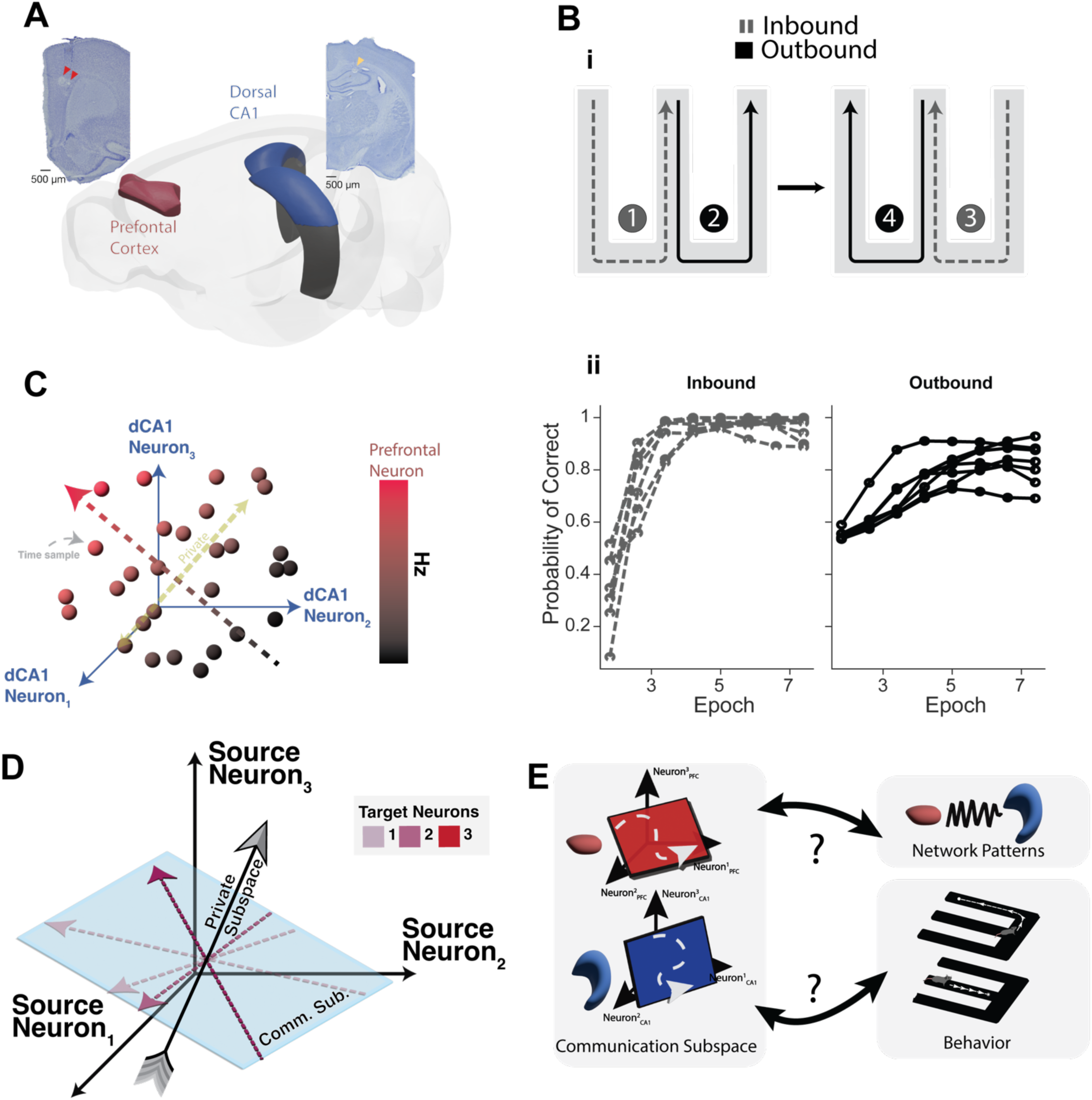
Connecting CA1-PFC Communication Subspaces to Memory-Guided Behavior and Network Dynamics. *(A)* Schematic view of the prefrontal cortex (PFC, highlighted in red) alongside dorsal CA1 (CA1, in blue). Insets display common electrode placements within the dorsal hippocampus (right) and prefrontal cortex (left). *(B)* Representation of the W-Track task learned over a single day, detailing the standard trajectory sequence numbered from 1-4. Dashed gray lines indicate inbound reference-dependent memory components, while outbound working memory phases are shown in solid black. During side arm visits, animals move to the center to receive a reward, whereas in the center arm, they recall their last visit, and choose the alternate side. In panel (ii), we show inbound performance of animals N=7 animals (left, gray dashed lines) and outbound performance (right, dark solid lines). *(C)* Demonstration of a one-dimensional communication subspace in a four-neuron system. Blue axes are formed by three CA1 source neurons. Scatter points display the firing correlation between the prefrontal target neuron and simultaneous CA1 neuron values. The bi-colored (red/black) dotted arrow represents the direction in CA1 space that promotes increased activity in the target neuron – termed ‘communication space’. The yellow dotted arrow indicates CA1 source directions with no impact on prefrontal firing denoted the ‘private space’. *(D)* Depiction of multi-dimensional communication within a three-neuron source and three-neuron target framework. Several target brain area neurons modulate activity spanning various dimensions of this subspace. ‘Private’ dimensions are those wherein alterations in source neuron activity do not cascade to downstream neurons. *(E)* Graphical representation of the study’s central questions. Two depicted activity spaces from the PFC (in red) and CA1 (in blue) are aligned with network patterns and behavior. The schematic of the communication subspace highlights core inquiries: How effectively do these subspace activities signal task behavior and network patterns? And to what extent do specific network patterns, such as theta coherence and sharp-wave ripples, influence the manifestation of activity in these spaces?

**Figure 1 – Supplement 1.**
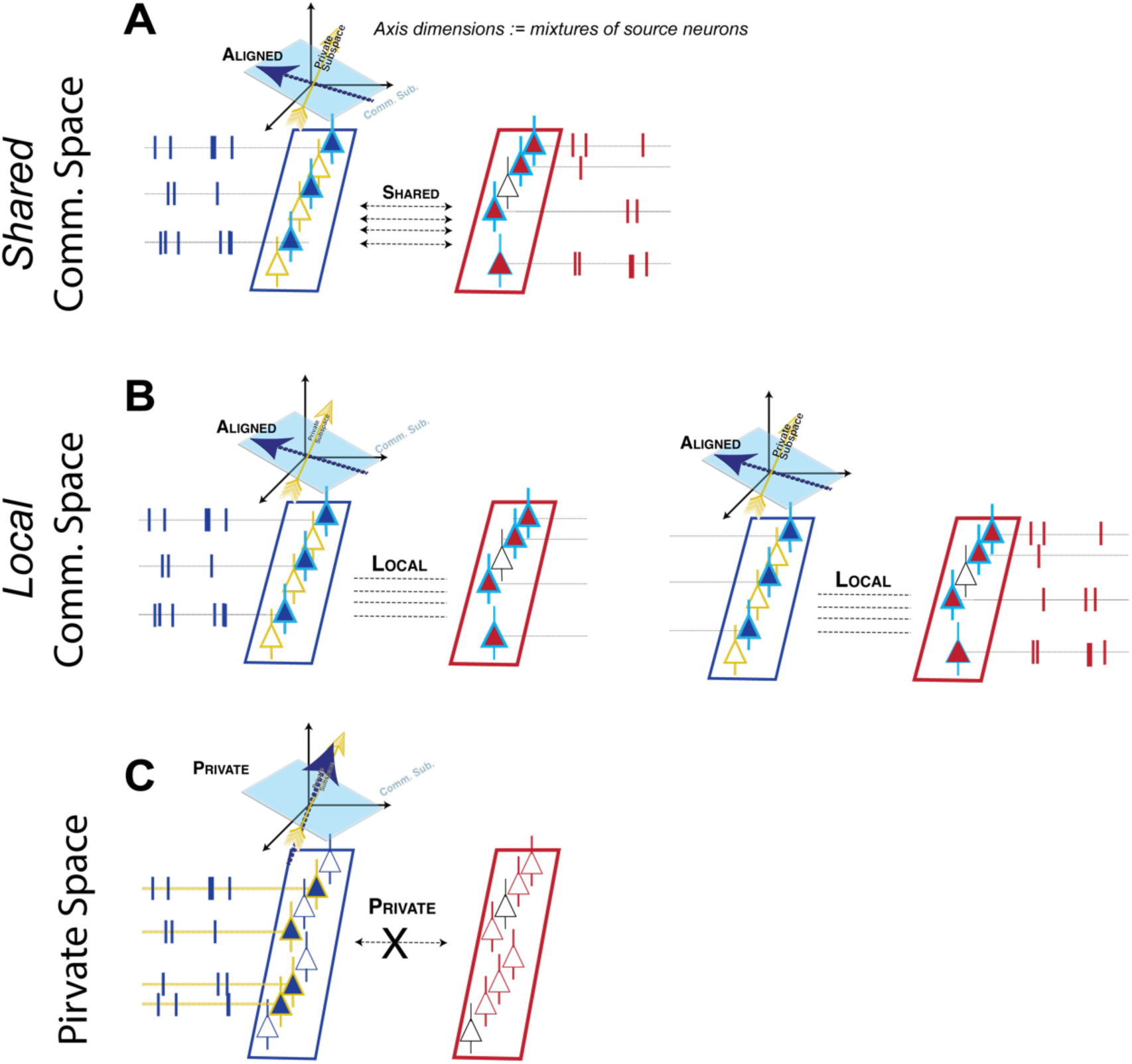
Schematic illustration of terminology. (A) Schematic shows an example of shared communication subspace activity (Veuthey et al. 2020) . Neurons which tend to cofire together across brain regions are depicted with a teal outline and neurons who do not statistically predict downstream neurons are depicted in yellow outline (private). When neurons who tend to cofire (communication space) actually do cofire, this is called “shared” activity. (B) We call neurons which tend to cofire, but fail to cofire in their typical activity ratio, local activity (Veuthey et al. 2020) – activity that aligns to communication space in either CA1 or PFC, but not both. In this sense, activity is not appropriately mirrored in the other area – i.e., unreciprocated. (C) The set of neurons which do not meaningfully add to extra-areal neuron firing rate prediction (Semedo et al. 2019) . These neuronal configurations differ from local, in that they avoid neurons that commonly cofire between brain areas. We did not identify this fraction in the current study, but it can be obtained by collecting neurons in reduced rank regression that do not contribute information.

**Figure 2.**
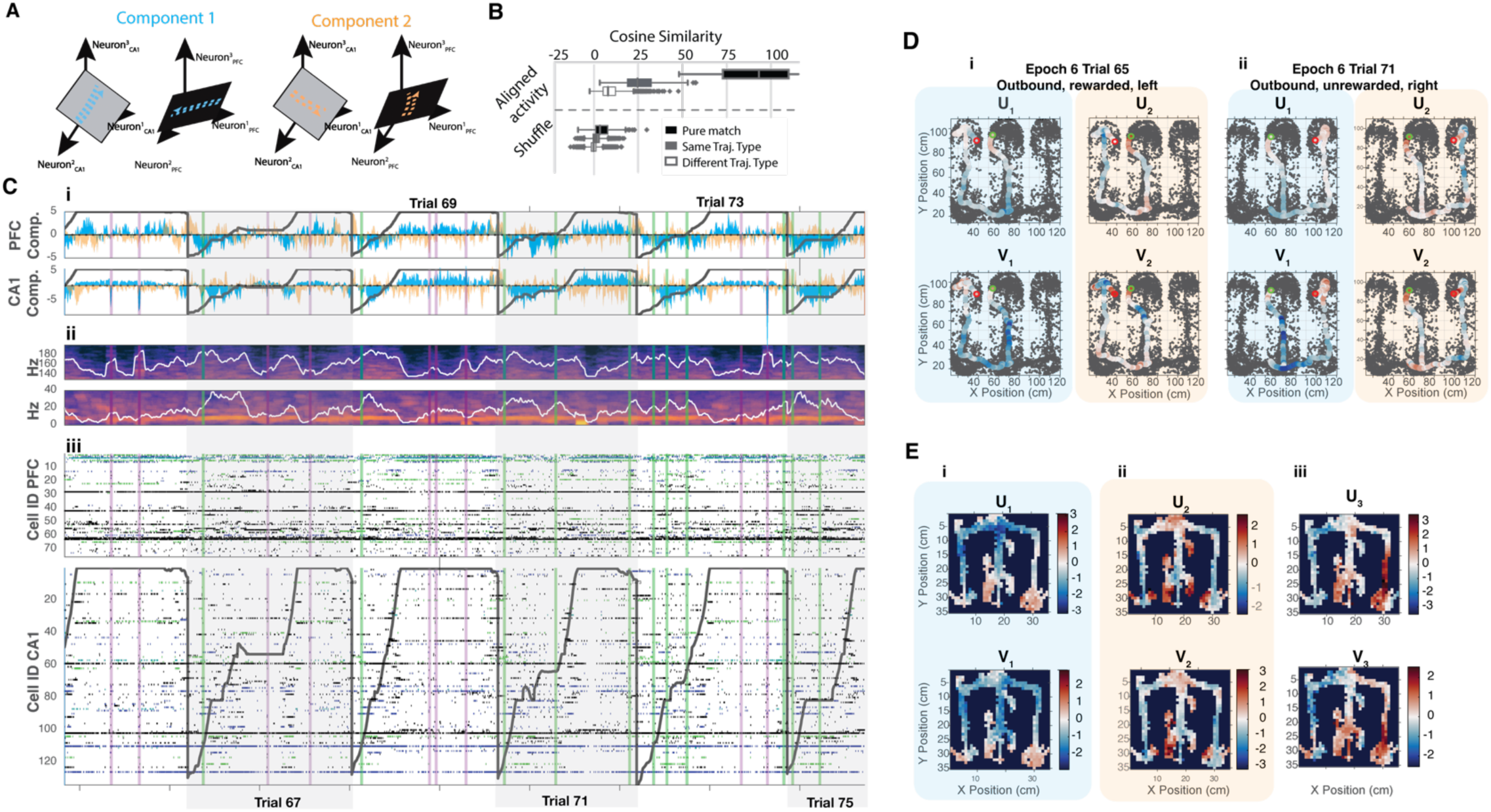
Canonical Activity Vectors in CA1 and PFC Display Consistent Coordination with Behavioral Outcomes in Animals. (A) Schematic representation of the first two subspace axes showing correlated neural activity in CA1 and PFC. Blue highlights the primary canonical variate of the communication subspace, while orange represents the second canonical variate activity axis. (B) The cosine similarity measurements indicate repeatability of canonical variate activity – how replicable canonical variate activity is across spatial trajectories. Each trajectory is binned into 100 locations and 10 aligned canonical variates for the 100 locations are cosine compared against other trajectories. Measurements are split by trajectories “Pure Match” indicates resampled points from the same exact trajectory (e.g. the 10^th^ outbound-left trajectory); “Same Traj. Type”, trajectories that match type (e.g. compare all within inbound-left or all within outbound-right); or “Different Traj. Type” trajectories that do not match type (compare inbound-left trajectories against inbound-right, etc). Cosine measurements are performed with 100 dimensional vectors of mean *UV* activity per resampled trajectory. (C)(i) Neural activity patterns along the principal correlated dimensions (1 in blue, 2 in orange) for hippocampal region (*U*) and prefrontal cortex (*V*). Dark grey trajectories represent the animal’s linearized path during consecutive outbound trials, illustrating repeatable communication structure. (ii), Spectrograms depicting theta and ripple frequency bands in the hippocampus. Levels of average band power are plotted in white, and periods of heightened theta (green) and ripple (pink) activities are highlighted via vertical shading (iii) Raster plots of spiking in CA1 and PFC. (D) Communication subspace activity components from (Ci) overlaid on the animal’s trajectory, color-coded from blue (indicating low activity) to red (indicating high activity). (E) Mean canonical variate activations across trajectories for an example epoch, revealing consistency with individual trial-based activations. i, ii, and iii depict the spatial means for *UV*_1_, *UV*_2_, and *UV*_3_ repsectively.

To sample the strongest time-varying activity in this communication space, we turned to a technique called Canonical Correlation Analysis (CCA) (Kim et al. 2023; Steinmetz et al. 2019; Veuthey et al. 2020). This technique, which exposes covarying activity patterns between two data sets, was used to identify major axes of interaction between the CA1 and PFC. This technique rotates the activity of two brain areas to expose their greatest covarying movements (**Figure 2B**). Many top components contain correlated activity, but far fewer components than neurons **(Figure 2 – Supplement 1)**. Here, we extracted the top two CCA canonical vectors to visualize their activity against the raw data. Each component has a hippocampal (*U*) portion and a prefrontal (*V*) portion that encodes how neurons that tend to covary in two regions activate, shown in **Figure 2B-D**. Indeed, the observed activities in two population directions are prominently covarying between CA1-PFC and repeatable across trials, with many outbound trials stitched together in **Figure 2C**. The top components exhibit consistent, repeatable deflections during sharp-wave ripples, during stillness periods in between ripples, and during running periods. These activations appear to reliably occur in positional space across trajectories shown in (**Fig. 2B**), and shown spatially in (**Figure 2C**), and appear to be repeatable across trajectories within an epoch (**Figure 2D**).

**Figure 2 – Supplement 1:**
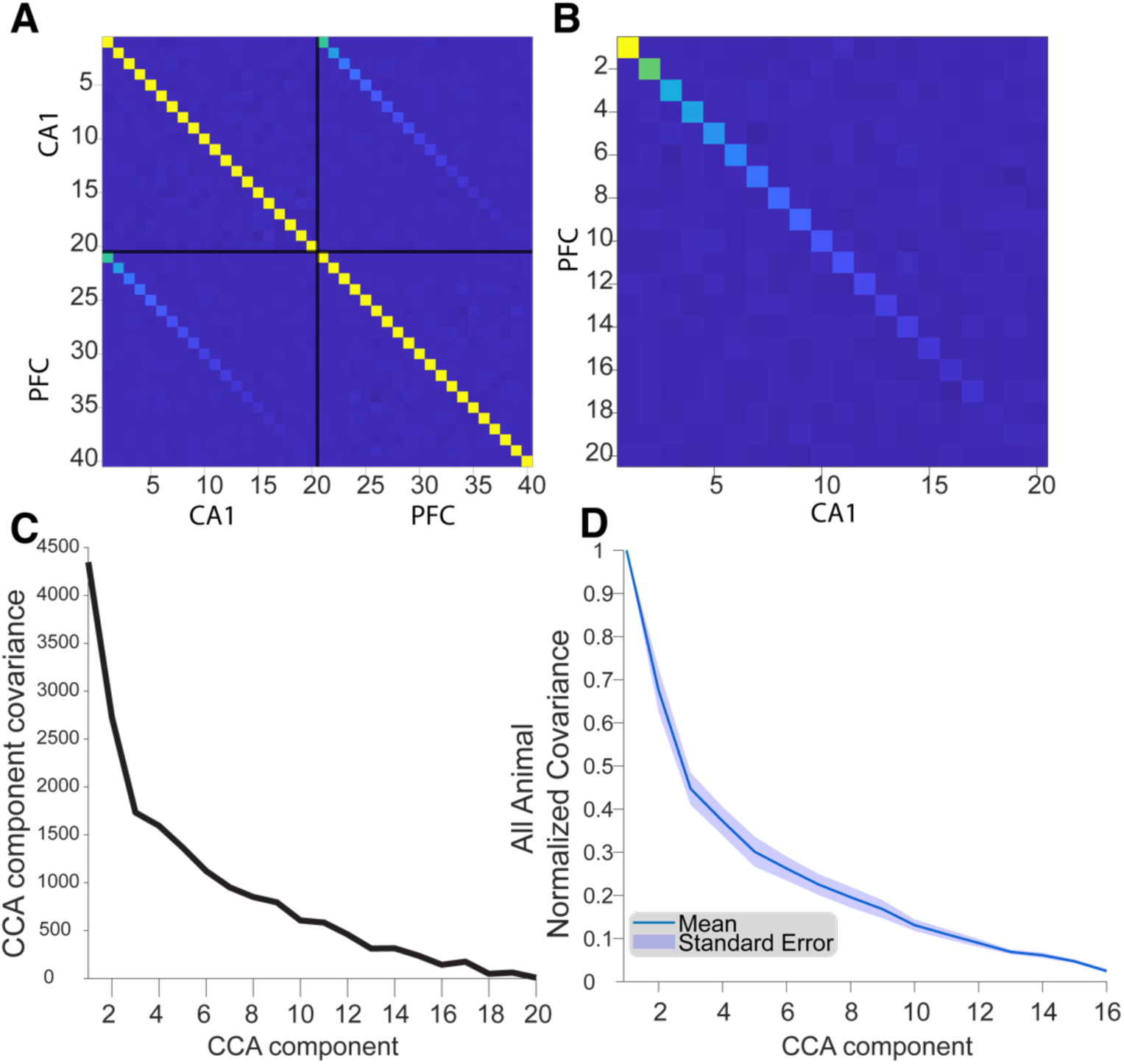
CCA Shared-component covariance. (A) Example covariance of U and V components over an animal session of data. This matrix is separated by black solid lines into CA1 and PFC components, with number of components equal the smallest rank of the two brain areas. Same brain area component correlations demonstrate orthogonalization of CCA components, whereas the CA1-PFC regions demonstrate the covariant overall relationship of a *U*_$_ to a *V*_$_component. (B) Like (A), except zoomed in along the upper-right CA1-PFC quadrant of (A). (C) A plot of the diagonal *U*_$_*V*_$_ covariance from one example animal, demonstrating rapid drop-off in covariance within the initial dimensions and exhibiting low-dimensional structure. The vast majority of the covariance occurs in the top 3 components. (D) Same as (C), except for summarizing all seven animals’ session data. Each animal was normalized such that maximal covariance was 1, where the solid blue line represents the mean and blue shaded area the standard error.

### Differential predictive performance and diversity of communication activity during rhythms

Next, we endeavored to understand and characterize the role of communication subspaces marked by rhythms. Given that we see a link between communication spaces and trajectory-specific position (**Figure 2B, E)**, and several studies support links between task behavior and rhythms (Haegens et al. 2014; Hyman et al. 2005; Iwata, Kiyonari, and Imai 2017; Moberly et al. 2018), we asked if there’s any evidence that communication subspaces can be changed or modified by rhythmic activity. We primarily focused on theta and ripple oscillations given previous reports of relationships to memory-guided behavior (Benchenane et al. 2010; Hasz and Redish 2020; Jadhav et al. 2012; Jones and Wilson 2005a; Joo and Frank 2018; Sigurdsson et al. 2010; Tang, Shin, and Jadhav 2021; Zhang et al. 2024). It is unclear how and if these states relate to the alignment of activity to the communication subspace and the implied shared activity conducted therefrom.

To characterize whether the communication space is stable or modulated in these rhythms, we defined periods of high and low rhythmic network patterns for each frequency band, the power or coherence **(Figure 3 – Supplement 1**, see also periods marked in **Figure 2**). We then sought to measure two key aspects within our windows: the dimensionality, which indexes the diversity of orthogonal population vectors predicting the target, and separately, the performance, measured by average cross-validated *R*^2^, which indexes the subspace’s ability to approximate information present in a source population about the target (schematized in **Figure 3A**). Dimension was measured at the point where adding a dimension induced no significant performance increase. An example is shown for CA1-CA1 and CA1-PFC interactions in **Figure 3B**, exhibiting lower between-area compared to within-area dimension as in Semedo et al. (2019). Like in Semedo et al. (2019), for each animal we re-partitioned source and target CA1 populations 50 times, both to control and match the number of target CA1 and target PFC cells, as well as to obtain a consensus of sample measurements that span all potential source CA1 neurons (**Figure 3C**). These source-target partitions were measured for each collection of windows per network pattern event.

**Figure 3 – Supplement 1:**
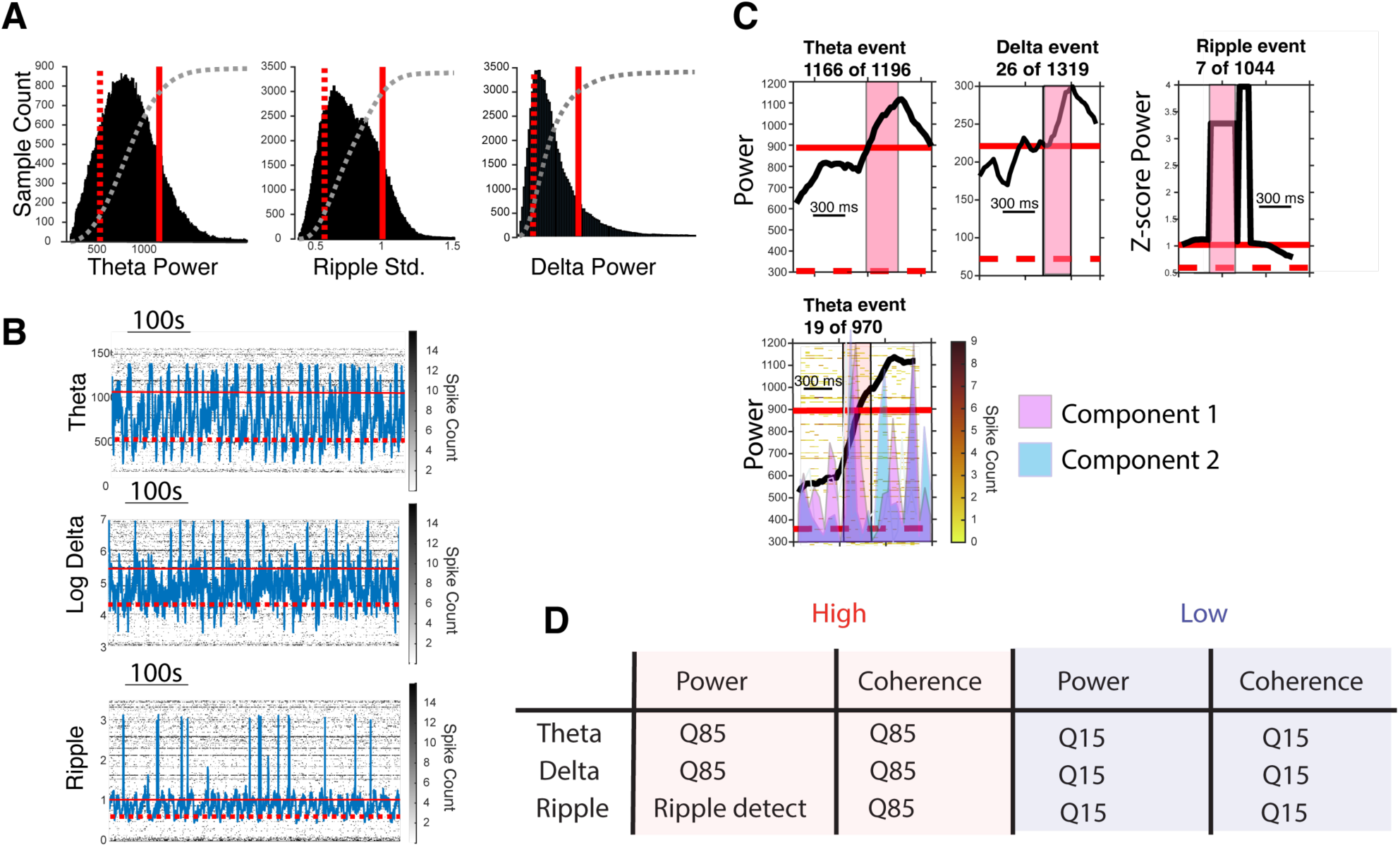
Windowing of high and low network pattern events. (A) Thresholding process for defining windows. High and low activity pattern windows are established using the 0.85 (high) and 0.15 (low) quantiles of the power distribution. Histograms of theta-, ripple-, and delta-band power are shown, with the upper quantile indicated by a solid red line and the lower quantile by a dashed red line. The light gray curve represents the cumulative distribution. (B) Example of window selection for three network patterns. The solid red line denotes the high network pattern threshold. When activity exceeds this threshold (solid red line), it becomes eligible for the selection of 300 ms sample windows (extending after the crossover point) – we sample the same number of windows per pattern type, and thus some windows will be discarded. Conversely, activity crossing the lower quantile threshold (dashed red line) becomes eligible for sampling 300 ms windows of low network pattern activity. (C) Examples of high events captured at crossover points are shown in the top three panels. The bottom panel displays an event with cell firing, where colored ticks represent spike counts. The color-shaded curves indicate activity matching the top two shared dimensions. Ripple curves were computed differently. When sharp-wave ripples were detected during quiescent bouts (see D), the curve was set to the global ripple mean power in standard deviations above the mean for its detected duration. Otherwise, the ripple curve was dynamically set to standard deviations above the mean. Ripple windows were therefore set by the onset of detection of ripples as previously described (Jadhav et al. 2012; Shin, Tang, and Jadhav 2019; Tang et al. 2017). (D) Tables specifying the details of window selection. Most network patterns utilize the lower quantile Q15 and upper quantile Q85 for defining low and high states, respectively. The exception is the sharp-wave ripple pattern, which uses Q15 for low sharp-wave ripple states but employs sharp-wave ripple event detection for high event selection. This is achieved by detecting ripple power at least 3 standard deviations above the mean, excluding times when the animal is moving (velocity > 4 cm/s).

**Figure 3.**
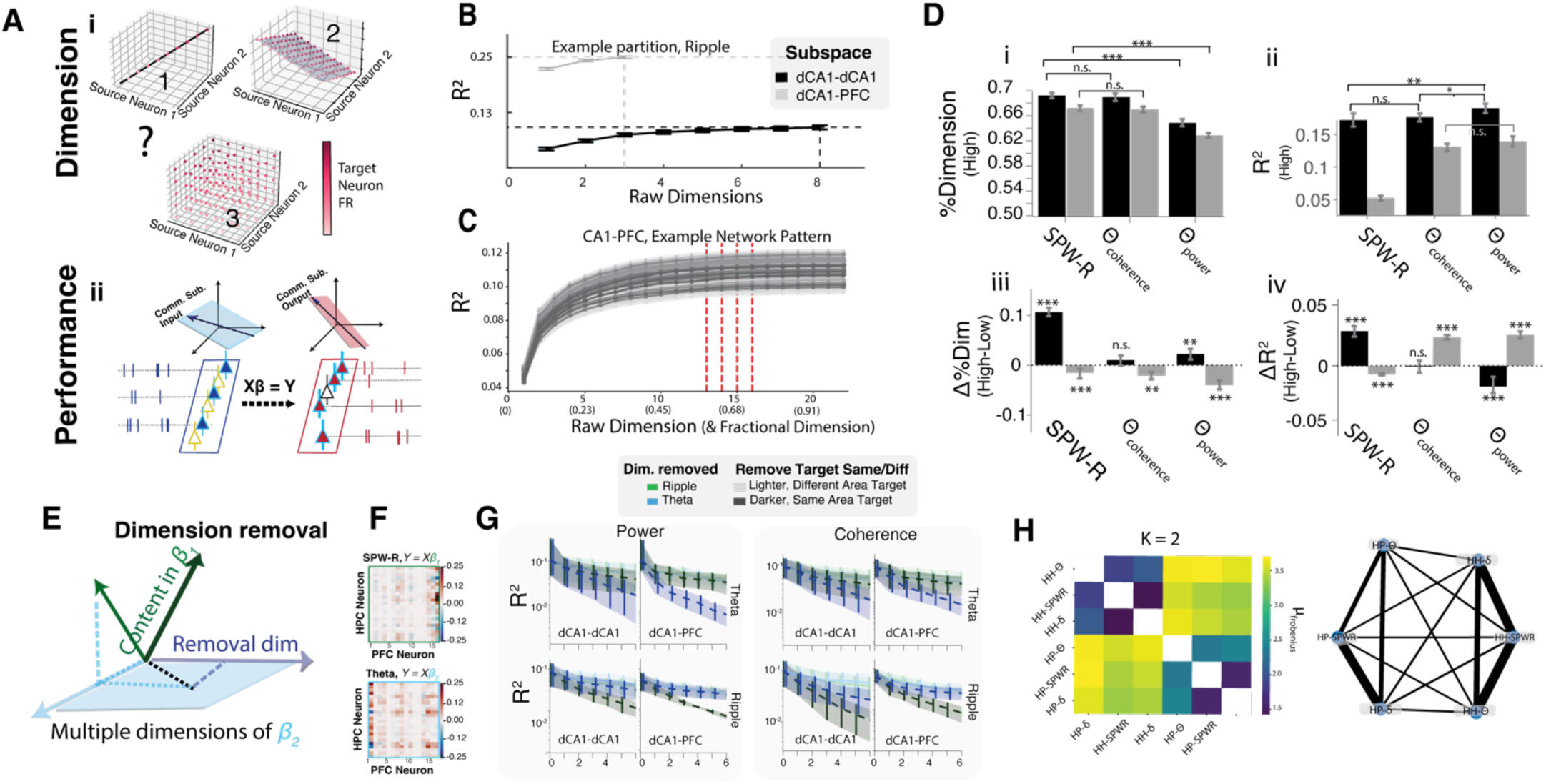
Assessing Dimensionality and Performance in Rhythm-Defined Communication. (A) Schematic illustrating the two primary aspects being measured from network patterns within or between areas: the dimensionality and performance of rhythm-defined subspaces. i, Dimensionality: The depicted subspaces represent possible communication dimensions within the neural data where target neuron(s) firing activity could potentially be influenced by various dimensionalities of source neurons. These three visualizations serve as exemplars, illustrating three potential dimensionalities to guide the reader’s understanding of the underlying principles and to preface what will be explored in sections B-D. ii, Performance: The graphic depicts a vector space defined by Beta coefficients. The blue vector space symbolizes the input aspect, originating from the firing rate of source neurons (CA1). This input is processed through a linear model, encapsulated by the Beta coefficient spaces, to project into the firing activity of target neurons. The output vector space is represented by target neurons, which can be located either in CA1 (the CA1-CA1 subspace) or PFC (the CA1-PFC subspace). (B) Measuring the Communication Dimension: This panel illustrates the process of assessing the communication dimension using a representative example. The y-axis depicts the model performance via coefficient of determination R^2^. A score of 1 represents the highest performance achievable with a full rank model, while values deviating from 1 indicate fraction of variance explained. As dimensions (shown on the x-axis) are incrementally added, performance is evaluated until it aligns within one standard deviation of the full-rank model’s performance (dashed horizontal lines: asymptotic performance, dashed vertical lines: optimal dimension). The dashed lines are decided when performance does not differ from the full rank model (within one SEM). In the depicted graph, intra-hippocampal CA1-CA1 dimension is marked in black, while the hippocampal-prefrontal communication dimension is shown in gray. (C) For each animal and respective network pattern, data is generated as outlined in the text, with dimensionality resampled as in (B) for every partition. X-axis, the integer dimension represents the number of neurons in the partition and the fraction represents the percentage of neurons used for asymptotic performance. Vertical red dashed lines indicate optimal dimensionality for 50 partition samples. (D) i, Comparison of dimension with network pattern type: CA1-CA1 dimension (black) and CA1-PFC dimension (gray). Error bars represent SEM. ii, Model performance plotted against high network pattern activity: C A1-CA1 dimension (black) and Communication dimension (gray). Error bars indicate SEM. iii, Dimensional difference from high to low network activity pattern. iv, Performance difference from high to low network activity pattern, with CA1-CA1 in black and CA1-PFC in gray. (E) Illustration of the dimension removal process, where a particular dimension from one communication space *β*_2_ (blue) is systematically extracted from content in the other *β*_1_(dark green) – see (F). The lighter green arrow represents content or a dimension in *β*_1_ after removing a dimension (gray line) from *β*_2_ (blue). (F) Two distinct Beta matrices, *β*_1_and *β*_2_, represent communication spaces 1 and 2, respectively. These matrices each delineate a relationship between CA1 neuron inputs and PFC neuron firing rates during SPW-R and Theta power rhythms. The color intensity within each matrix conveys the strength of the neuronal connection. (G) Outcomes of successive dimension removal are graphically detailed in terms of performance, quantified as the percent variance (y-axis) predicted after dimension removal (x-axis), with shading of 95% confidence intervals The most pronounced decline, seen when dimensions from a subspace’s native rhythm are removed, highlights its critical role in neural communication. Similar gradients and interval overlap indicate the shared information between the two spaces, with the divergence points underscoring the distinctiveness of each space’s informational content. (H) A heatmap, constructed using the Frobenius norm of the differences between the communication spaces 1 and 2, captures their relative similarity or distinction. The varying color intensities spotlight disparities between the spaces, while the network diagram elucidates the interplay and strength of relationships among different neural oscillatory patterns.

Figure 3D shows the results of this process. Our comprehensive analysis of rhythmic communication patterns yields interesting insights, suggesting that not all rhythms express a communication subspace equally. As previously seen, one tends to find diminished dimensions for inter-areal communication (black, CA1-CA1, Figure 3D (i)) compared with intra-areal (gray, CA1-PFC, Figure 3D **Figure 3 – Supplement 2**) for each type of rhythm-defined window (Figure 3D **i**, two-sample t-test, p < 0.001 for each, N=350 partitions, 50 per animal, FDR corrected, Q=0.001). This finding aligns with previous work (Semedo et al. 2019; Srinath et al. 2021) by showing reduced inter-areal communication dimensions for all rhythms when comparing high intra- vs inter-areal states. Notably, however, we make a novel observation that the presence of network patterns (high state minus low state) strengthens this difference in dimensionality (Figure 3D **iii**; **Figure 3 – Supplement 2**).

Beyond these similarities, network patterns also differ in their effects on the communication dimension. We observed a significant reduction in the diversity of both CA1-CA1 and CA1-PFC dimension diversity during high theta power states (two-sample t-test, p < 0.001, N=350 partitions, 50 per animal) relative to high SPW-R and high theta coherence states (Figure 3D i). Interestingly, the presence of theta coherence does not significantly modulate CA1-CA1 communication dimensions (p = 2.67e-1), while the presence of SPW-R and theta power does. However, all three high states demonstrated reduced CA1-PFC dimension compared to low (Figure 3D **iii**).

Apart from the diversity and dimension of network patterns, we can also investigate how these patterns influenced the predictability: how well is target region activity predicted from the source. Not surprisingly, CA1 cells better predict other CA1 cells than PFC cells (Figure 3D **ii**; **Figure 3 – Supplement 2**) with significantly higher *R*^2^ values during all three states (p < 0.001, all three rhythms). On the other hand, CA1-PFC rank regression prediction was maximal for theta-based patterns, and the presence of theta patterns (high-low) increased CA1-PFC regression prediction (Figure 3D iv). In contrast, sharp-wave ripples strongly enhanced CA1-CA1 prediction. Overall, theta rhythms were associated with enhanced periods of predictable, lower-dimensional inter-areal communication. Meanwhile, SPW-Rs exhibited enhanced prediction more intra-regionally, with overall higher CA1-CA1 and CA1-PFC subspace diversity.

**Figure 3 – Supplement 2:**
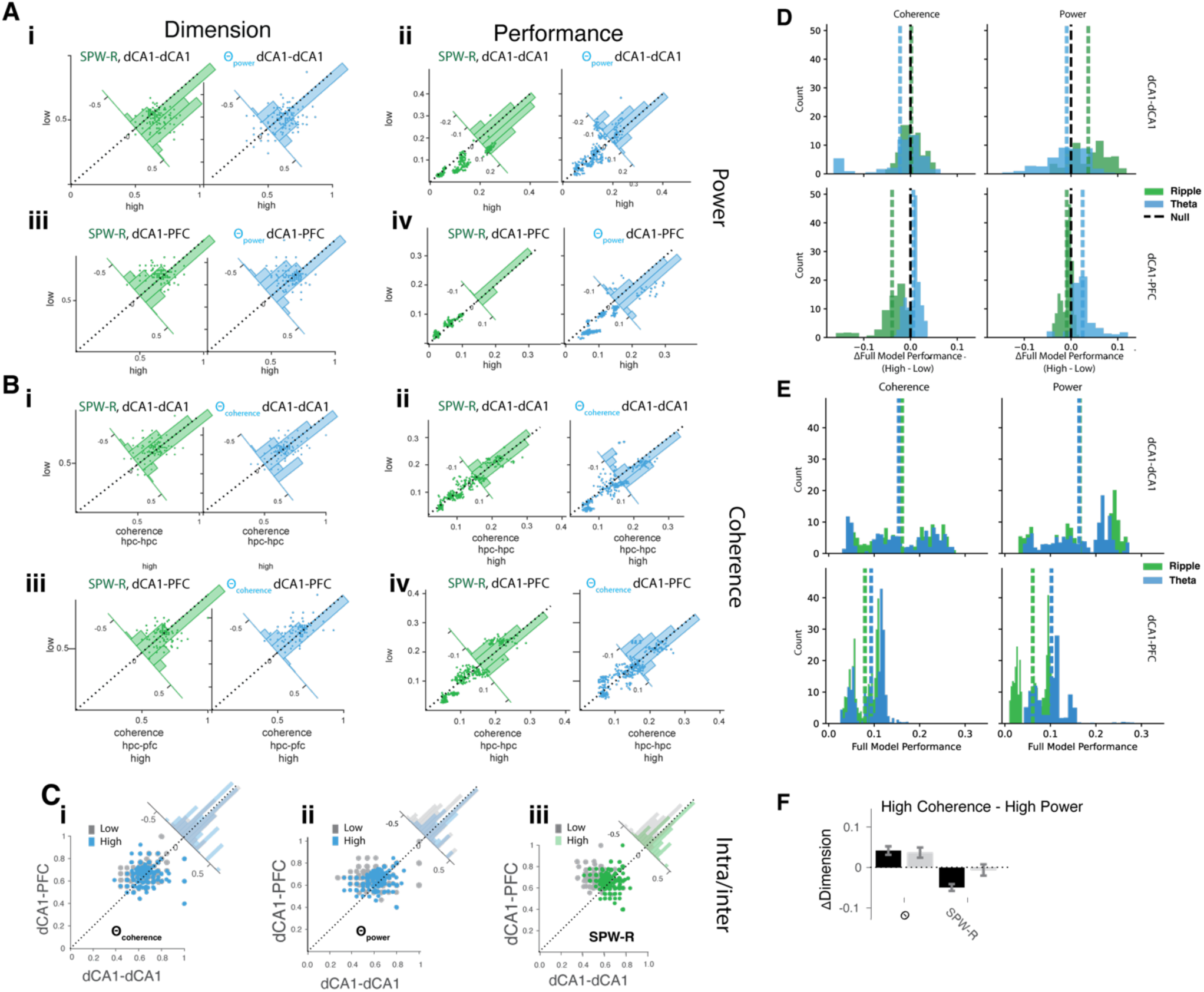
Additional Insights into Rhythm-Defined Communication. (A) Each subplot in A and B depicts measurements for all 50 partitions per animal. Panels split by those measuring dimension in the left column versus performance in the right. Each subplot contains an overlaid inset histogram showing the difference for that partition measured during high and low activity. A broad asymmetric lean of the histogram suggests a difference between high and low activity patterns. (B) Same as A, except for coherence. (C) Each panel measures the fractional dimensionality for intra-hippocampal versus inter-regional communication for each of the 50 partitions of 7 animals. Colored points indicate measurements for high activity for a given rhythm and gray for low activity for a given region. i, ii, and iii cover windows of theta coherence, theta power, and SPW-R, respectively. Insets display the differences between the x- and y-axis for high (colored) and low (gray) activities, respectively. (D) Change in performance from high to low pattern activities for each rhythm. The first column shows coherent activity, the second for power; the first row is CA1-CA1, and the second is CA1-PFC. Dotted lines indicate distribution means. (E) Same as (D), but for full-model performance. (F) The bar plot summarizes differences in fractional dimension between high coherence and high power, 95% CI given by error bars.

Given that rhythmic network patterns (theta and SWP-Rs) coincide with differences in the communication subspaces — different statistically but sometimes reasonably similar in dimension or performance, we sought to grasp the commonality between network pattern communication. To what degree do the key neuronal population vectors overlap? To investigate this, we sequentially removed overlapping dimensions (Figure 3E) between subspaces from different rhythmic patterns as described in Semedo et al. (2019). Removing each overlapping dimension and remeasuring the performance allows an estimate of the degree of entanglement between subspaces. Figure 3E shows an example of two subspaces that were previously partitioned as depicted in Figure 3A. Using a dimension of predictive content in space *β*_1_, if we project the population vector into another space, we can find concomitant key dimensions of space *β*_2_, and subtract them from the content, thus removing them (Figure 3E). This conceptual notion is implemented by a method that achieves this effect (Semedo et al. 2019) (see Methods).

Examining the result, we note that the initial dimensions of subspaces tend to be most entangled, indicated by the uniform performance drop when removing a space’s own dimensions versus a non-matching space’s dimensions (Figure 3F). For all spaces, removing their own dimension causes the quickest descent. The more unique the content, the fewer cumulative dimension removals before its performance differed significantly from separating others. Notably, the intra-areal CA1 spaces overlapped most, and CA1-PFC spaces diverged much more quickly. The separation between communication spaces was more pronounced when comparing high vs low power states rather than coherence states. Subspace differences for theta and ripple power emerged within 3-4 dimensions, while more dimensions were needed to observe differences between theta coherence spaces (Figure 3G). This finding suggests there is greater separation between the communication subspaces during high vs low theta power states and ripple power states, compared to theta coherence states.

To confirm this, we also computed distances between *β*_1_ and *β*_2_. Namely we resampled the Frobenius norm ∥ *β*_1_ – *β*_2_ ∥*_F_* of the two spaces repeatedly permuting the positions of target neurons (Figure 3H). This measures the typical total Euclidean distance between two population vectors in *β*_1_ and *β*_2_, as well as the sum of squares of the difference matrix’s singular values. Furthermore, this method demonstrates that the initial dimensions (specifically the first two) most prominently distinguish between inter- and intra-areal components in the neural code, with rhythms following suit as the next most important feature. And notably, the more quiescent ripple and delta periods show stronger connections to each other than to theta.

### Theta coherence is not a dominant feature of the overall communication space and declines with learning

We found it interesting that theta coherence exhibited no measurable performance gain in prediction controlled against high theta power (Figure 3D). Previous studies either suggest CA1-PFC theta coherence to be periods of increased CA1-PFC spiking correlation (Benchenane et al. 2010) during choice periods, putatively for memory retrieval (Jones and Wilson 2005b; O’Neill, Gordon, and Sigurdsson 2013; Sigurdsson et al. 2010) or memory encoding (Battaglia, 2011). Some even suggest that theta coherence has no consequence on interareal spike correlations (Bygrave et al. 2019; Nardin et al. 2023). We therefore used the CCA method, which provides a temporal readout of shared/local communication without filtering window times. This method allows us to test whether theta coherence is temporally associated with communication along the top subspace components without the filtering window paradigm, which merely samples high and low states. And further, if so, does coupling between coherence and communication appear at a particular point within learning?

We first characterized how theta coherence dynamics are modulated over learning in this task. To examine how coherence changes over learning epochs, we utilized two methods of coherence measurement: multi-taper coherence and weighted phase lag index (WPLI; Vinck et al. 2012; Mitra and Bokil, 2007). Both showed strong low-frequency bands, most prominently the theta band (Figure 4B). We found that overall coherence on the track dropped significantly over the learning period for all spatial positions, rather than gaining steadily over time (Figure 4C). We confirmed this general decline in coherence over learning epochs by z-score normalizing and min-max normalizing each animal’s coherence and WPLI values (Figure 4C).

Because different task phases or regions of the W-track may coincide with differing cognitive operations, e.g. coherence-mediated recall, we separated these measurements over space and trajectory type (Figure 4D). Spatially-binned coherence over epochs strongly confirmed a learning effect. Coherence began high in nearly all areas of the maze—most prominently in the center arm where animals presumably engage in working-memory-guided decision-making. Moreover, coherence elevated throughout the entire outbound trial. However, over the course of the eight learning sessions, coherence declined. During outbound journeys, coherence remained above average levels, while during inbound returns, coherence was suppressed to around average. Overall, coherence declines from its peak well before animals hit peak performance at epoch 5-6 (Fig. 4C**-D**). This declining pattern of coherence over learning epochs implies a role for coherence in online learning or encoding of the task rather than memory retrieval, aligning with prior theories (Battaglia et al. 2011).

We next turned to characterize the fluctuations of the top CCA components with respect to coherence. Significantly, the range of canonical variate values reflecting shared activity substantially varied between high and low theta power states, which did not occur for unreciprocated, local activity. However, these same values of shared activity did not vary as much with different theta coherence states (Figure 4F). Since this relationship may change over epochs, in a more refined manner, we trained a linear model once per epoch using the top shared communication components to predict theta coherence. Over all the animals, theta coherence exhibited a low relation with aligned communication (*R* = 0.03), declining over time (Figure 4 **G, H**). In contrast, all animals exhibited a strong relationship with theta power (*R* = 0.25) (Figure 4 **G, H**), with several individual animals showing very strong relationships. This theta power relationship grows over learning epochs, asymptoting around the time that place fields and population vector codes stabilize (Tang et al. 2021; Zielinski, Shin, and Jadhav 2021).] Notably, our animals exhibit no trends in mean velocity (Figure 4C, ANOVA, p=0.14; Mann-Kendall, p = 0.90). Acceleration, on the other hand, exhibits a minor trend (Figure 4C, Mann-Kendall, τ=0.64 p=0.03) over learning across animals, though we did not find individual epoch acceleration to be well-differentiated across epochs (Figure 4C, ANOVA, p=0.65).

**Figure 4.**
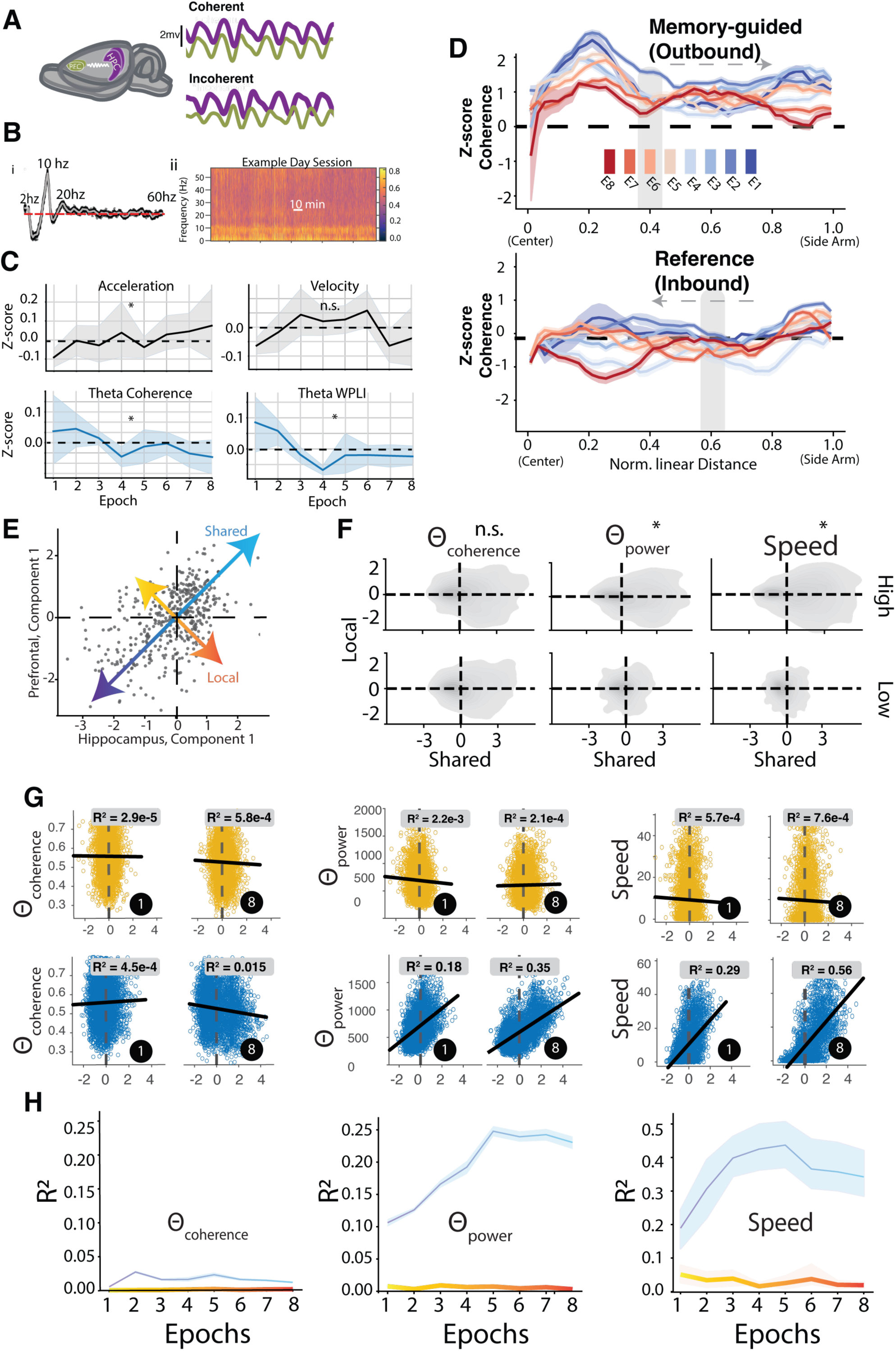
Change in shared activity correlations with theta power, and not theta coherence, over the course of learning. (A) The purple (CA1) and green (PFC) schematized traces are local field potentials filtered in the theta band. Coherent theta activity holds a relatively constant phase lag relative to incoherent activity. (B) Panel i presents a mean coherence spectrum, highlighting prominent peaks in both delta and theta power ranges. Panel ii offers a snapshot of an entire session, showcasing the prevalence and interplay of strong theta and delta coherence throughout. (C) Panels depict epoch averages of theta coherence, theta WPLI, velocity, and acceleration, each normalized by z-scoring session values per animal. Each solid line represents mean values and shading the 95% confidence intervals for N=7 animals. Notably, acceleration significantly increases (Mann-Kendall, p=0.03; ANOVA, p=0.64), while theta coherence significantly decreases for both multi-taper (Mann-Kendall: p=0.03; ANOVA, p=0.33) and WPLI (Mann-Kendall, p=0.03; ANOVA, p < 0.005) theta coherence. Velocity exhibits no significant changes (Mann-Kendall, p=0.15; ANOVA p=0.09). (D) Comparative analysis of coherence against W-track linear distances, categorized into outbound (top panel) and inbound (lower panel) trajectories. The displayed curves, shaded for clarity using 95% confidence intervals, represent mean z-scored coherence values. A discernible decrease in coherence is evident across most track areas over time. (E) Example of shared versus local activity in Figure 3. (F) Kernel density plots showing the distribution of samples for windows of high versus low activity for theta power, theta coherence and speed. The densities indicate that the range of observed values in shared and local activity depends more strongly on power than coherence. * Indicates p-value < 0.001 via permutation test of the univariate shared axis distribution. (G) Scatter plots of shared/local components for an example animal from the first CCA component (x-axis) versus theta power/coherence/movement speed (y-axis) levels. Samples are taken for all times. Coefficient of determination is displayed atop each subplot. Top row depicts local components (yellow), bottom shared components (blue). (H) Summary of the coefficient of determination values found for (N=7) all animals. Shaded region depicts 95% confidence interval. The coherence values tend to peak early and at a much lower value than the asymptotic strength of the CCA relation with theta power.

In summary, we found that theta coherence, in fact, declined over learning epochs rather than increased, with coherence being highest early in learning across spatial areas of the track. Additionally, theta coherence exhibited minimal relation to the shared CA1-PFC communication space, while theta power showed a stronger relationship. This suggests that theta power and not coherence is a major feature of the functional communication CA1-PFC subspace during this memory task.

### Shared communication subspaces but not local subspaces encode task-relevant behavior

Finally, we sought to better understand whether these fluctuations encode task-relevant behavior. To that aim, we adopted and trained a simple linear model, resampled, and retrained repeatedly with Markov Chain Monte Carlo techniques. We used two activity categories related to the top 10 canonical CA1-PFC variates in the communication space: shared and local activity. More conceptually, aligned activity refers to source and target neural movements that are reciprocated and coincide with one another. In contrast, local activity describes a situation where either the source or target has varied in the communication space without commensurate activity in the other area. This has been referred to by Veuthey et al. (2020) as “local activity” and is sometimes indicative of future communication but is not yet a shared fluctuation. Figure A displays the activity of CA1 and PFC projected onto the top canonical variate. The *U* (x-axis) encodes how much CA1 activity matches the communication covariate vector and the *V* (y-axis) encodes how much PFC matches the same covariate. If we look at the projection onto the unity line, this is where changes (or motions) of neural activity in one brain area reciprocated into the other brain area – i.e. shared activity. The orthogonal axis to the shared motion shows unreciprocated motions, i.e., local subspace: positive or negative imply population vector motions along one area’s communication axis without concomitant expected motions in the other brain areas.

**Figure 5.**
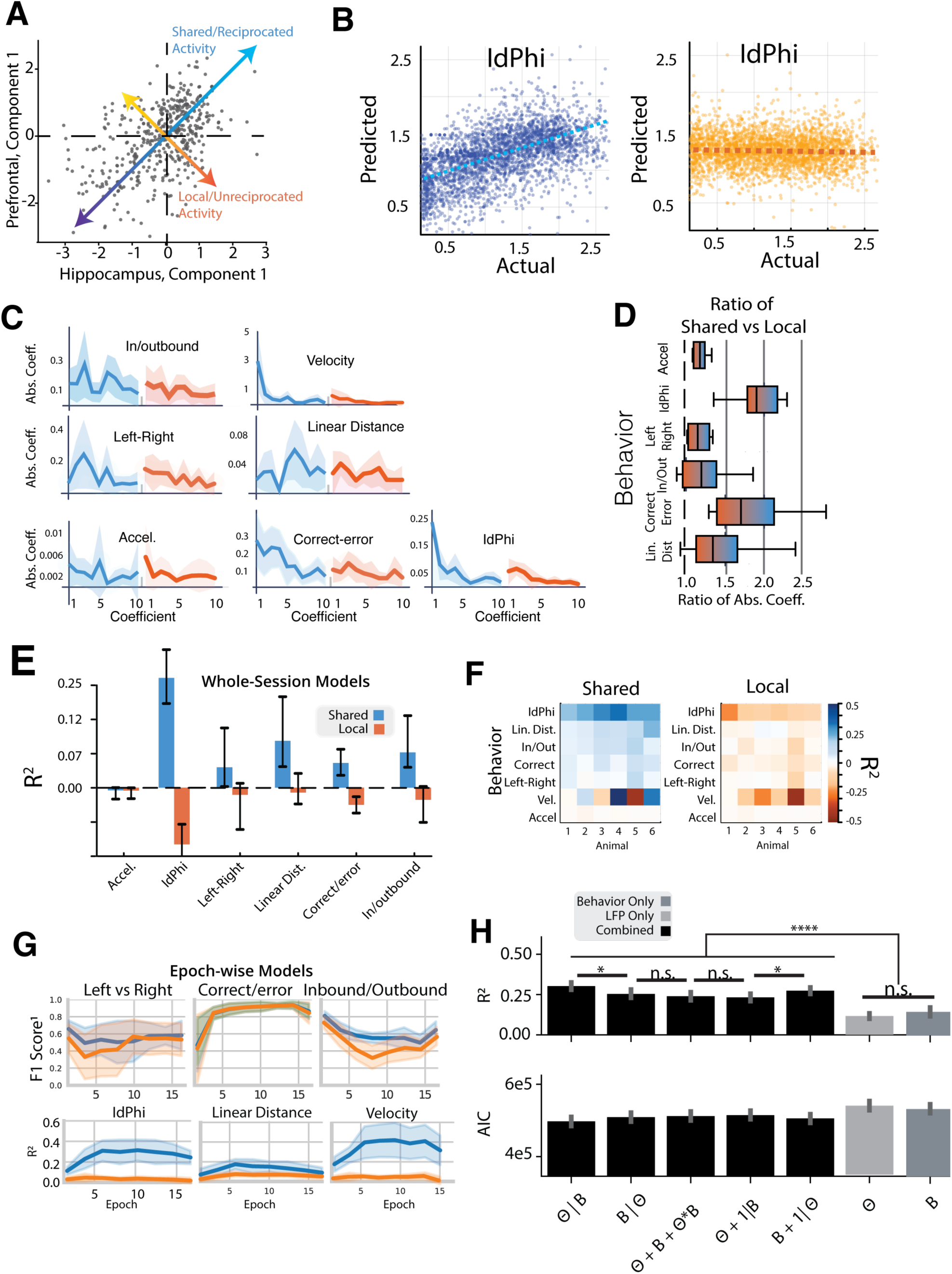
Shared CA1-PFC Activity Predicts Task Behavior Better than Local Activity. (A) A scatter of time samples on the U (CA1) and V (PFC) axes. Blue arrow illustrates aligned activity between CA1 and PFC, while the orange arrow indicates unreciprocated CA1 activity with corresponding PFC movements. (B) Outcomes from the Bayesian linear model predicting IdPhi, a metric indicating vicarious trial and error. Analysis contrasts reciprocated (blue) vs. unreciprocated (orange) movements. (C) Display of linear model coefficients corresponding to seven distinct behaviors on the W-track task. Coefficients are color-coded: shared (blue) and local(orange). Coefficients, resampled via Markov chains, come with shaded 95% confidence intervals. (D) A depiction of the ratio between shared and local cumulative coefficient magnitudes. A unity reference (1) is marked by a dashed black line, representing equal contributions. (E) *R*^2^ values (y-values) for each linear model predicting a behavior (x-axis) from canonical variates. Shared activities significantly predict task-relevant behaviors above zero, while local activities do not. (F) A breakdown of mean *R*^2^ values as in (E) for individual animals, categorized by aligned (left panel) and orthogonal (right panel) components. (G) Performance of models trained over individual single-day learning epochs. The bottom panel shows *R*^2^ values for continuous variables over epochs, while the top panels display F1 scores for classification variables. Shared components (blue) tend to outperform local components (orange) in predicting task-relevant behaviors, except correct/error. Shaded regions represent 95% confidence intervals. (H) Comparison of models predicting the top CCA shared component from various combinations of theta power (θ) and continuous behavior (*B*) information. Models are specified using R-like formula notation, where hierarchical terms are indicated by the vertical bar operator (|) and interaction terms by the asterisk operator (*). For each model, the top value represents the *R*^2^, and the bottom value represents the Akaike Information Criterion (AIC), which accounts for model complexity and goodness of fit. Models incorporating both behavior and theta power information consistently exhibit better performance in terms of higher goodness of fit *R*^2^ and lower AIC values compared to models using only one source of information. Error bars depict 95% confidence intervals.

We sought to understand if activity along several possible shared and local activity axes explains behavior. Thus, we desired to train a linear decoder to predict behavior from the top K shared or local activity components. CCA exhibits as many canonical covariates as the smallest neuron count (the minimal rank of the brain areas), but in practice, far fewer contain significant shared activity. Most animals express less than 10 significant axes, and therefore, we used the top 10 covariates aligned/reciprocal activity and orthogonal/non-reciprocal activity to train a linear decoder to predict task behavior. Many task-relevant behaviors could be approximated with this very crude model (N=6 animals), but only using the reciprocated communication activity (Figure 5B**, E, F**). Task-relevant variables include the phase of the task (outbound/inbound), linear distance along the track, IdPhi (integrated local head direction changes), the turn direction, and the trial’s correct versus error state. Key among these, IdPhi (Figure 5B) measures an animal’s integrated head deflections locally in time and has previously been used to index vicarious trial-and-error (VTE) and deliberative decision behavior (Papale et al. 2012; Redish 2016; Santos-Pata and Verschure 2018; Schmidt et al. 2013); this index of VTE was highly represented in shared CA1-PFC activity across animals. Such predictive patterns are replicated across animals (Figure 5F).

In addition to this task-relevant aligned encoding, we also examined the unreciprocated, local communication (Figure 5, *orange*). In every case, surprisingly, the local subspace’s unreciprocated activity provided little to no predictive power. This can be seen in the flat response of predicted IdPhi versus actual (Figure 5B) as well as the poor determinacy over task variables in Figure 5E**, F**. Linear models trained to predict behavior with local activity produced muted beta coefficients for local components (Figure 5C) relative to reciprocated components (Figure 5D). These findings collectively imply that while shared dynamics between CA1 and PFC significantly predict behavior, the individual subspace activity within the regions does not, highlighting the unique behavioral significance of communication space driven shared activity.

To examine whether learning changed this behavioral prediction, we turned to training GLM models in an epoch-wise fashion (Figure 5G). We created a series of cross-validated models within each epoch trained on the top 10 CCA components predicting six of the behaviors. Continuous behaviors, such as IdPhi, linear distance, and velocity, exhibited enhanced predictive performance *R*^2^ in the shared versus local components, which rose over the learning period and stabilized in the well-learned period. For the classification variables, local and shared produced more similar predictions across epochs as measured by F scores. Only the correct-error classifier increased over epochs, and only the in/outbound classifier produced a period of significant difference.

Given communication strongly indicated behavior (Figure 5) and our previous analysis showed increasing correlation of the top CCA component with theta (Figure 4), we asked whether the local field potential effects could be entirely explained by behavior, vice-versa, or required both. Perhaps behavioral state determined local field potentials, and the former differences (Figure 3**, 4**) could be explained solely by behavior. To this end, we created a series of models to explore the effect of behavior alone, theta alone, or models that combined information from both sources predicting the contiguous behaviors discussed above (speed, IdPhi, and linear distance). Notably, behavior and theta power generated similar performance for the top CCA component when pooling the prediction of velocity, linear distance, and IdPhi prediction (**Figure 5H**). But to our surprise, all models that combined both sources of information created better predictions with lower AIC (t-test2, p < 0.01 for all interaction versus univariate; Figure 5H). The improved predictive power from interactions suggests that behavior and rhythms, while correlated, have independent influences on communication subspace activity.

**Figure 6.**
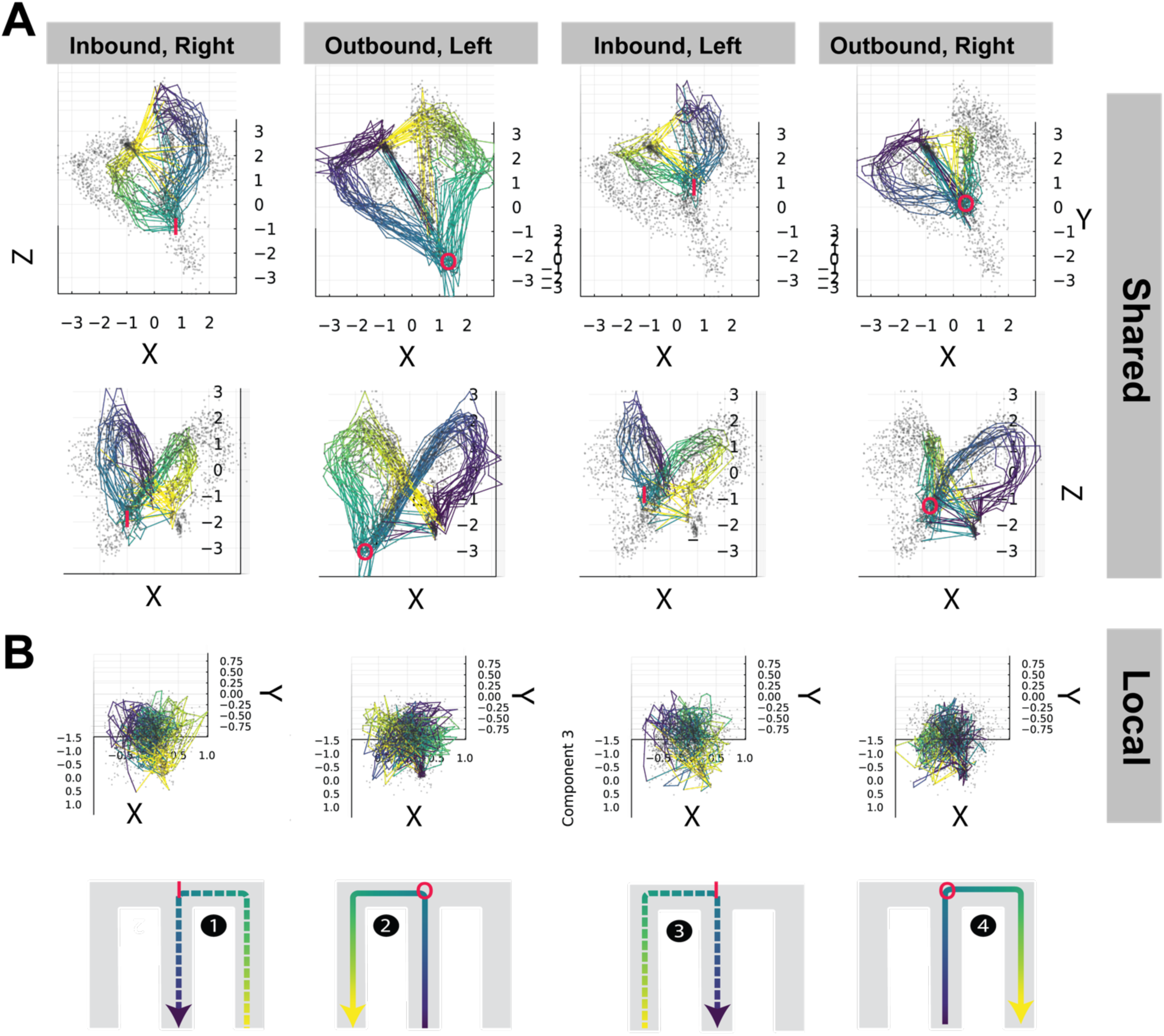
The communication subspace organizes trajectories in nested ring attractors. (A) Combined animal communication shared subspace. Trajectories of each type were ordered, binned, and concatenated across animals to help visualize whether the communication subspace organizes activity into manifold and whether that manifold organizes behavior. In this case, we color-labeled the manifold by the linear distance bins (center=violet, side-arm=yellow), positions shown in the panels below (B). The shared communication activity organizes trajectories into rings, with remote-like bisection events between choice and reward locations. Each trajectory is colored in accord with panels below (B), while other trajectory points are depicted by small gray scatter points. The choice zone is demarcated by a red ‘o’ symbol for outbound and ‘I’ for inbound. This choice locus differs clearly across the many samples of each trajectory type and, together with other spatial differences, explains the ability of linear decoders to sense behavior in this space. (B) This structure is mostly absent for local communication activity, activity matching the communication subspace but not coordinated across the two areas. A small amount of positional clustering can be observed.

**Figure 6 – Supplement 1.**
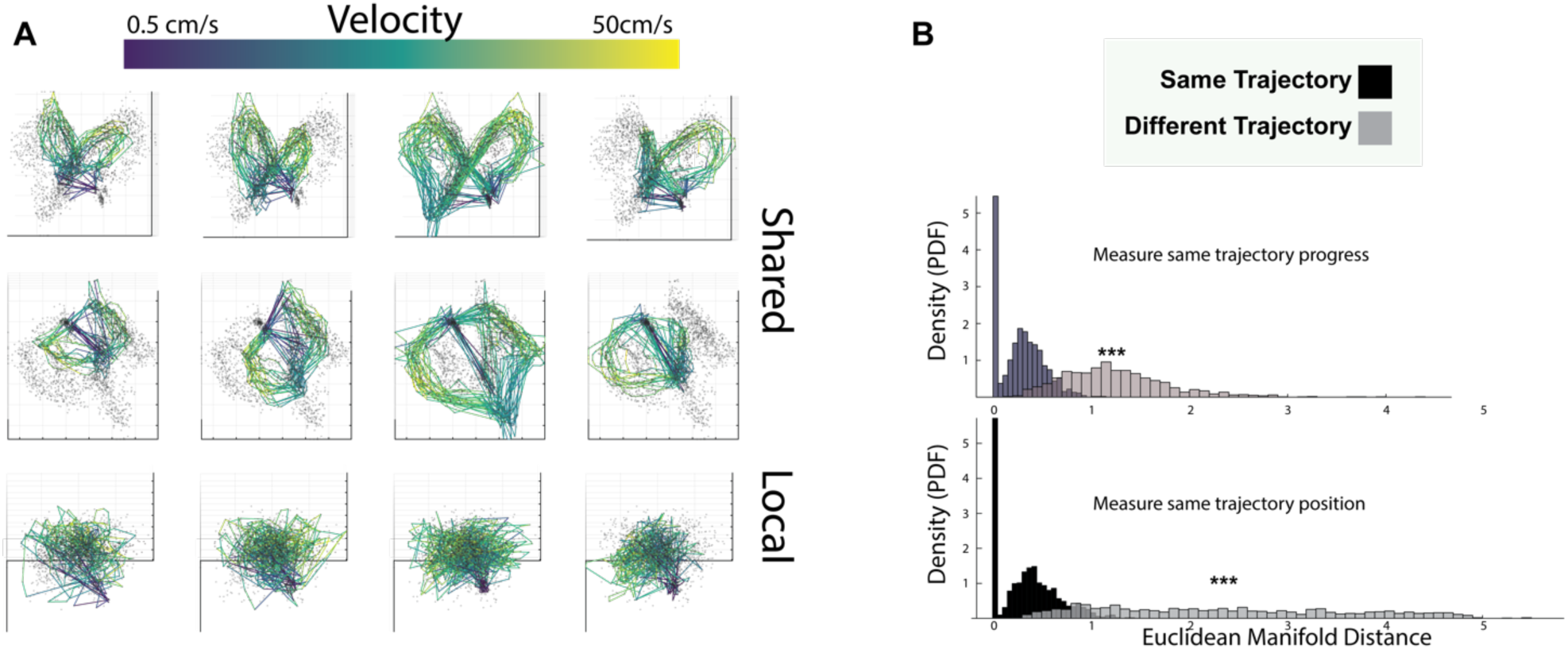
Velocity and manifold distance quantification. (A) This shows the same manifold as in **Figure 6AB** but colored by the average animal velocity. Ring mainfold bisection events can be seen as lower velocity states. (B) Probability density normalized histograms of manifold Euclidean distance within (black) and between (black) communication manifolds, where distances are taken for samples sharing a linear distance.

Lastly, given that communication subspace activity well-explained behavior, we wondered whether the subspace manifold structure itself appeared to structure behavior into an orderly manifold. To this aim, we created a pooled dataset across animals based on trajectory type (Shin et al. 2023; Tang, Shin, and Jadhav 2023), and then we color-coded the CA1-PFC shared and CA1-PFC local activity manifold by the associated mean behavioral labels (Figure 6A-B**; Supplemental Video 1**). This revealed the presence of differential cyclic structures for shared activity manifolds -- structures not apparent for local activity manifolds. Each shared activity manifold possesses a unique cyclical path proceeding clockwise through communication space (looking towards the midline crease) Nested ring sizes signaled trajectory type (inbound/outbound) and left versus right. This was confirmed by measuring the difference in Euclidean distance between points sharing linear distance within and between communication manifolds (Wilcoxon-Rank Sum p < 0.001, between vs. within distance 95% CI 1.90 – 2.06); **Figure 6 – Supplement 1B**). Taken together with the clockwise observation, points on different rings are neighboring not by linear distance from the center – but rather along trajectory progress, with rings folded at the choice and reward points. Furthermore, robust separation occurs at the choice point between manifolds

These rings interestingly exhibit bisection. The activity at the outer wells (center and sidearm) jumps to the other side of the ring closer to the choice point. From a neural geometry perspective, these points (reward wells and choice points) are closer to each other on the third axis, where the rings are folded downwards, increasing their similarity and signaling, potentially reducing the energy landscape barrier for neurons in the communication space moving between these points. We also examined this space labeled by average velocity. Notably, the activity bisecting between the reward arm and choice regions is associated with regions of lower velocity (Figure 6 **– Supplement 1; Supplemental Video 2**). In summary, labeling the shared activity manifold by speed and space makes it apparent *why* these spaces may produce behavioral prediction, even with simple linear decoders.

**Figure 6 – Supplement 2.**
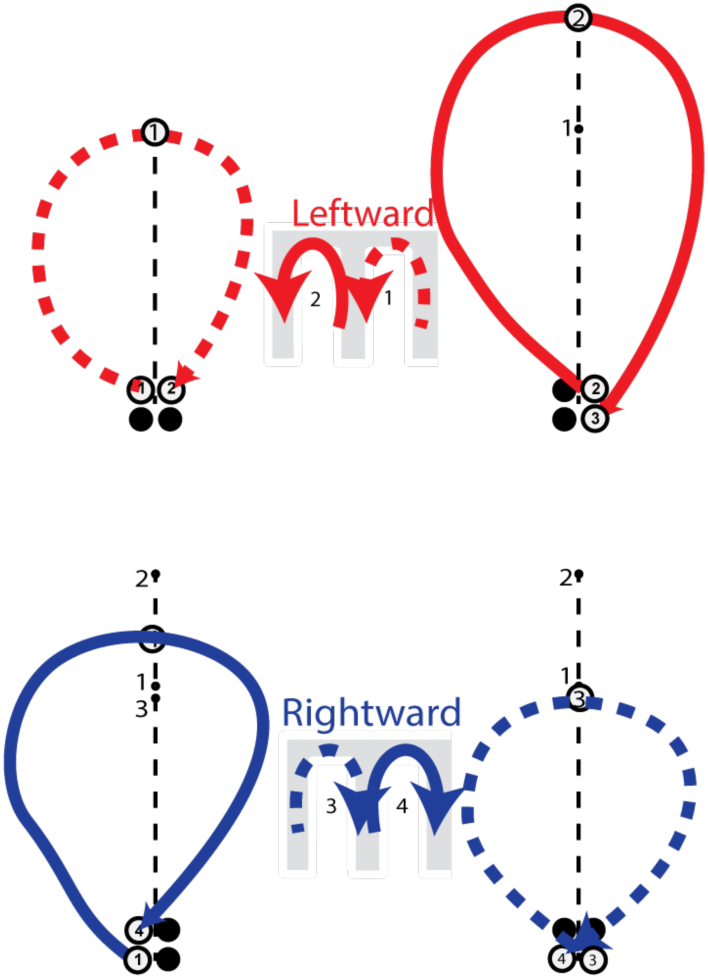
Attractor Hypothesis. The four trajectories are schematized. Inbound is shown in dashed line style, outbound in solid line style, leftward is red, and rightward is blue. The trajectories form neighboring rings—all proceeding clockwise (Figure 6). Different trajectories are labeled 1-4 near their respective choice point schematics. Active start/end and choice points are shown in white. Though we have not explored the initial positions, we hypothesize that neighboring ring attractors could distinguish trajectory class and may terminate near the start of the upcoming trajectory. In this manner, such CA1-PFC coactivity could implement a form of task logic in the communication space.

**Table S1.** Cell Counts. Cell counts per animal within CA1 and PFC.

| Animal | CA1 | PFC |
| --- | --- | --- |
| 1 | 60 | 30 |
| 2 | 44 | 26 |
| 3 | 44 | 16 |
| 4 | 39 | 21 |
| 5 | 41 | 48 |
| 6 | 34 | 38 |
| 7 | 132 | 75 |

## DISCUSSION

Our examination of hippocampal subspace dynamics during memory-guided behavior yields several key insights. We found that shared CA1-PFC activity within sparse communication subspaces predicts task behavior (**Figure 4-5**). Our results suggest that this is because CA1-PFC shared activity forms organized ring-like manifolds arranging various behavioral states (trajectories) in the W-track task. Further, dimensionality reduction techniques revealed that prominent rhythms modulate subspace properties (Figure 3). Notably, theta periods exhibited increased predictability of CA1-PFC activity and reduced CA1-PFC communication dimensionality, compared to greater intra-CA1 diversity during SWP-R events (Figure 3). Surprisingly, we found that theta coherence did not coordinate shared subspace activity as well as theta power. Theta coherence actually declined over learning and exhibited far less shared activity coupling, in contrast to an increasing prominence of theta power shared-activity coupling over the course of learning (Figure 4).

Memory-guided behavior requires hippocampal-prefrontal interactions (Floresco et al. 1997; Maharjan et al. 2018; Rezayat et al. 2022; Shin and Jadhav 2016). Our approach sought to measure activity across networks via shared communication subspace versus local activity via private communication subspaces. (Figure 2**, 5, 6**). This approach directly suggests that shared information is better related to immediate behavior compared to locally held information. This does not preclude such local activity from having delayed effects on memory-guided behavior; indeed, Veuthey et al. (2020) found that local, unreciprocated activity often precedes changes in shared activity by up to a second and frequently follows changes during early learning in the motor cortex. In a similar vein, we saw that the local manifolds appear rather disorganized, though future studies may find greater order when examining local spaces against time-lagged behavior; these spaces may contain prospective or retrospective information yet to be coordinated across areas. An example of this time-lagged interaction occurs in Kaufman et al. 2014, Elsayed et al. 2016, and Veuthey et al. 2020.

We have shown here, on the other hand, that shared space well-organizes the animal’s task, track, and velocity space. These structures contain potential clues for how CA1 and PFC networks coordinate to implement the W-track task rule. The ring-like manifolds emerging out of the shared activity differ by task phase **(Figure 6 – Supplement 1)**, but potentially relate to geometric structures seen in these areas individually (Tang et al. 2023) . Notably, the neural activity in the communication space indexes the current location and trajectory subtype (**Figure 6 – Supplement 2A**). Neighboring trajectory loops could act like attractors – paths that neural activity may bias to remain near based on the shared coactivity. And progress coding may bear some resemblance to that noted in prefrontal cortex by El-Gaby et al. 2023, who found cells indexing trajectory progress in prefrontal cortex. Though in our study, such information emerges in CA1-PFC coordination.

Differences also emerged at the intra-hippocampal level between theta and ripple states. Intra-hippocampally, our results revealed differences in diversity (Figure 3), with ripples exhibiting a larger increase in diversity. One potential explanation is that distinct neuron populations activate within these rhythms, such as CCK+ and OL-M cells during theta power versus PV+ cells during SPW-Rs (Klausberger et al. 2003; Somogyi et al. 2014). Differential neuronal inhibition levels likely contribute to the differences in spreading intra-hippocampal activity observed during ripples and theta states. Much like in the case of the visual cortex (Semedo et al. 2019), we note that the dimensionality changes in more subtle than drastic ways. The overall changes tend to be on the scale of 10% differences in assembly composition. Likewise, the shared spaces have similar magnitudes of predictability, from 5-20%.

Prior studies posit theta coherence aids in retrieval processes in alternation tasks. However, these results on the W-track does not appear to support that view. Prior results also failed to find a difference in theta coherence between correct and error trials (Tang et al. 2021). A more likely explanation resembles older theories that propose theta coherence is conducive for plasticity and synaptic weight changes during early learning, and the theta rhythm parcellates ideal windows for inducing LTP and LTD (Buzsáki and Moser 2013; Dragoi and Buzsáki 2006; Hyman et al. 2003; Wilson, Varela, and Remondes 2015). This can allow theta coherence with another region to take advantage of this window for linking assemblies of ongoing experiences. Thereafter, ripple associated reactivation can stabilize weakly linked traces (Benchenane, Tiesinga, and Battaglia 2011). This hypothesis also explains the increased diversity of patterns in pre-consolidated inter-areal theta coherence and ripples (Figure 3) and the reduced correlation to a stable cross-area response (Figure 4).

These findings suggest that theta coherence, if any role at all, is more likely to play a role in early learning or encoding rather than a retrieval process of extant memory. The network pattern of coherence is elevated early during learning and diminishes over time, similar to sharp-wave ripples. This pattern could support the encoding model proposed by Benchenane et al. (2011) more than the models suggesting pure working memory retrieval interactions between CA1 and PFC, such as those proposed by Griffin and Hallock (2013).

Our study contains some key limitations. The analyses were restricted to only awake brain states. Possibly, important population vectors and local/shared interaction balances substantially differ between sleep and wake. Examining communication architectures during sleep presents an important future direction for fully characterizing hippocampal subspace dynamics. Additionally, prior work has found limited impact of nonlinearities on the dimensionality and geometry of visual cortical communication subspaces. However, using residual rather than total activity helps remove nonlinearity. The W-track task, however, lacks the regular trial structure permitting residual removal previously used. So, we focused on full z-scored spiking, which subtracts a mean. Total activity previously yielded a similar result (Semedo et al. 2019). Still, nonlinearities can alter linear approximation in nuanced ways.

Lastly, we characterized the effects of rhythms, behavior, and changing spike alignments on a stable, session-wide subspace, which has pros and cons. The pro observes how activity aligns with an overall fixed state of CA1-PFC coordination. The con is an inability to examine acute but stable coordination structures in different epochs, especially early epochs. Future studies could utilize a temporally windowed approach to understand how temporary communication dimensions arise and either stably incorporate them into the already stable CA1-PFC structure or fail to do so. Separate epoch analysis may reveal learning-related changes in the rhythmic modulation and behavioral structure of these shared CA1-PFC spaces.

One difficulty is that our recordings contained limited numbers of certain events like ripples, presenting analysis challenges. Using lower quantile thresholds to split by epoch could help overcome limited events like ripples for analysis. This task also involved two dissociable rules with likely differential working and reference memory demands. Comparing subspace modulation by rule type presents another interesting direction, including testing the specificity of theta’s growing communication influence. More broadly, future work could manipulate rhythms to causally probe their impacts on dimensionality. Physiological identification of neuron types in the subspaces could reveal a cause for warping or angular changes. Much remains unknown regarding how subspace topography sculpts routing, presenting exciting frontiers for mapping this structure-function landscape.

These results provide evidence for the importance of reciprocal interaction on behavior and the influence of rhythm on communication subspaces, particularly theta power, not coherence. Together, these observations advance an understanding of how oscillatory states, memory-guided behavior, and spiking-based communication motifs entangle during the routing of information through neural populations.

## ACKNOWLEDGEMENTS

This work was supported by the National Institutes of Health (R01MH112661) to S.P.J. and a T90 training grant (5 T90 DA 32435-7).

## AUTHOR CONTRIBUTIONS

S.P.J. provided project oversight, manuscript writing, and literature review, secured funding, and contributed to editing, reviewing, and conceptual framework. R.A.Y. contributed to manuscript writing, literature review, experimental design, data analysis, and conceptual framework, and was involved in editing and reviewing the manuscript. J.D.S. was responsible for data preprocessing and data collection. Z.G. assisted with the literature review and data analysis.

## METHODS

### Subjects

Seven adult male Long-Evans rats served as subjects in this study. Their weights ranged from 450-550g at the ages of 4-6 months. The Institutional Animal Care and Use Committee at Brandeis University approved all procedures, which followed the United States National Institutes of Health guidelines. Data from all rats were previously reported in earlier work (Tang et al. 2021; Zielinski, Shin, and Jadhav 2019).

### Animal Pre-training

The animals were habituated through daily handling for several weeks prior to training. After habituation finished, the rats underwent food deprivation to reach 85-90% of their *ad libitum* weight. The rats were then pre-trained to run along a linear track (∼1m long) to receive rewards of sweetened evaporated milk. They also became accustomed to spending time in a high-walled, opaque sleep box (∼30 x 30 cm), as described previously (Jadhav et al. 2012; Tang et al. 2017; Zielinski et al. 2019). Following pre-training, the animals received surgical implantation of a multi-tetrode drive, with LFPs measured relative to a cerebellar ground screw. Electrodes were not moved for at least 4 hours pre- and during recording (Tang et al. 2021).

### Microdrive Implantation

Surgical implantation procedures were as previously described (Jadhav et al. 2012; Tang et al. 2017, 2021). Briefly, six animals of seven animals were implanted with a multi-tetrode drive containing 32 independently moveable tetrodes targeting right dorsal hippocampal region CA1 (−3.6 mm AP and 2.2 mm ML) and right PFC (+3.0 mm AP and 0.7 mm ML) (16 tetrodes in CA1 and 16 in PFC for four animals; 13 in CA1 and 19 in PFC for two animals). One animal was implanted with a multi-tetrode drive containing 64 independently moveable tetrodes targeting the bilateral CA1 of dorsal hippocampus (−3.6 mm AP and ±2.2 mm ML) and PFC (+3.0 mm AP and ±0.7 mm ML) (30 tetrodes in CA1 and 34 tetrodes in PFC). On the days following surgery, hippocampal tetrodes were gradually advanced to the desired depths with characteristic EEG patterns (sharp wave polarity, theta modulation) and neural firing patterns as previously described (Jadhav et al. 2012; Tang et al. 2021; Zielinski et al. 2019). One tetrode in corpus callosum served as hippocampal reference (CA1 REF), and another tetrode in overlying cortical regions with no spiking signal served as prefrontal reference (PFC REF). The reference tetrodes reported voltage relative to a ground (GND) screw installed in skull overlying cerebellum. Electrodes were not moved at least 4 hours before and during the recording day (Tang et al. 2021).

### Behavioral Training

Following recovery from surgical implantation (∼7-8 days), the animals were food-deprived again and pre-trained on a linear track for at least 2 days. Then the novel W-track sessions began (Figure 6A; ∼80 x 80 cm with ∼7 cm wide tracks). The rats were introduced to the W-track for the first time and learned the task rules over eight behavioral epochs on a single day. Each 15–20-minute epoch was followed by a 20–30-minute rest session in the sleep box (total recording time ∼6 hrs). As depicted in Figure 6A, the rats performed a hippocampus- and prefrontal-dependent continuous alternation task on the W-maze (Jadhav et al. 2012; Maharjan et al. 2018; Zielinski et al. 2019). The rats had to alternate their visits between the left (L), right (R), and center (C) wells to receive automated rewards upon crossing infrared beams at the reward sites. The four correct trajectory types were: C-to-L, L-to-C, C-to-R, and R-to-C. Learning performance was estimated via a state-space model (Jadhav et al. 2012, 2012; Zielinski et al. 2019).

### Electrophysiological Recording

The data analyzed here were partially reported previously (Tang et al. 2021, 2023; Zielinski et al. 2019). The hippocampal and prefrontal cortex activities were recorded using 30-64 tetrodes positioned in the right dorsal hippocampal CA1 region and prefrontal cortex. Recording sites were verified via histology. The individually movable tetrodes targeted the desired regions. Data were collected with a SpikeGadgets acquisition system (Tang et al. 2017; Zielinski et al. 2019). Spike signals were bandpass filtered 600 Hz to 6 kHz and sampled at 30 kHz. Local field potentials were bandpass filtered 0.5 Hz to 400 Hz and sampled at 1.5 kHz. The overhead color CCD camera tracked animal’s position and running speed via affixed LEDs (30 fps). Single units were identified through manual clustering based on amplitude, principal components, and spike width using MatClust software (M.P. Karlsson), as described before (Tang et al. 2017; Zielinski et al. 2019). Only well-isolated neurons with stable waveforms were included (Zielinski et al. 2019). Neurons firing below 0.1 spikes/second on average were excluded across all analyses.

### LFP Preprocessing and Coherence Quantification

The local field potential (LFP) underwent bandpass filtering in the delta (0.5–4 Hz), theta (6–12 Hz), and ripple (150–250 Hz) bands. Zero phase infinite impulse response (IIR) Butterworth filters were utilized. Coherence calculations were performed with the Chronux (http://chronux.org/, RRID:SCR_005547) MATLAB software package for spectral analysis (Zielinski et al. 2020).

### Rank Regression Analysis

#### Partitioning

Spikes were counted in 50ms bins across the run recording periods. To examine related activity between the two areas, we reasoned that fluctuations in hippocampal neurons could relate to fluctuations in prefrontal cortex neurons and vice versa.

The hippocampal and prefrontal neurons were matched by their average firing rates. The distribution of firing rates was divided into 20 equal-range bins. The minimum number of neurons from each bin was randomly selected from both areas. Since more hippocampal cells were typically recorded, the distribution of selected target cells matched the prefrontal distribution. The target population size ranged from 11 to 65 units (mean: 29.2). Partitioning was repeated 50 times, selecting different sets of source neurons and mean-matched target neurons each time. The partitions were consistent across different methods of generating windows of interest (described below) to control dimensionality analyses. Each source and target pair underwent reduced rank regression (Semedo et al. 2019).

#### Windowing of Rhythmic Periods

High local field potential (LFP) activity windows were selected when the oscillation of interest exceeded 85% of the band power, using a 300ms window beginning at the threshold trigger of the oscillation. Low oscillatory activity windows occurred when power fell below 15% of the band (*SFig. Window Partitions*). Overlapping windows were removed, both within and between different activity patterns. The windows were matched so that all oscillation patterns and strengths had approximately uniform windows across the recording period.

Statistical tests based on comparisons of window states (high/low, intra/interareal, rhythmic band) were corrected by Benjamini-Hoch for multiple comparisons.

#### Rank Regression

We first correlated spiking activities between source and target regions using linear regression:

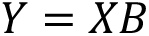

*X* represents the *p* source neurons by *T* timepoints matrix, where *T* timepoints come from *N* windows each containing *t_w_* timepoints. *Y* represents the *T* timepoints by *q* target neurons matrix. The coefficient matrix *B* (*p* × *q*) combines activity in each row of *X* to minimize the squared error. We optimized the solution using ridge regression:

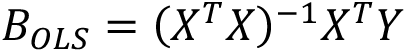

To test whether a subspace of the source neurons could still predict the target activities, we applied reduced rank regression. This gradually took the first *m* principal components of *B_OLS_* to reduce its rank to *B_RRR_:*

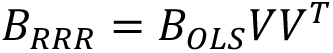

We then re-predicted the target population with the new *B_RRR_* matrix to find the smallest number of dimensions where performance plateaued within 1 standard error of using all neurons. Cross-validation determined this optimal subspace dimension.

#### Removal of Predictive Dimensions

We first correlated spiking activities between source and target regions using linear regression:

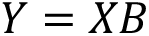

*X* represents the *p* source neurons by *T* timepoints matrix, where *T* timepoints come from *N* windows each containing *t_w_* timepoints. *Y* represents the *T* timepoints by *q* target neurons matrix. The coefficient matrix *B* (*p* × *q*) combines activity in each row of *X* to minimize the squared error. We optimized the solution using ridge regression:

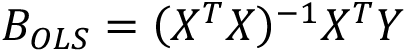

To test whether a subspace of the source neurons could still predict the target activities, we applied reduced rank regression. This gradually took the first *m* principal components of *B_OLS_* to reduce its rank to *B_RRR_*:

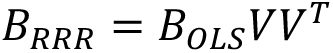

We then re-predicted the target population with the new *B_RRR_* matrix to find the smallest number of dimensions where performance plateaued within 1 standard error of using all neurons. Cross-validation determined this optimal subspace dimension.

To examine the effects of removing source population activities along predictive dimensions, we first calculated the activity fluctuation:

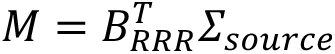

where Σ*_source_* is the covariance of source activities, and *M* represents fluctuation along each predictive and source dimension. Taking the SVD of *M* = *UDV*^T^, where *U* is source activity fluctuation along predictive dimensions.

We calculated *Q* by gradually removing leading columns of *U*, starting from the first *u_m_* columns down to just the *u_m_*th column, where *m* is the previously found optimal subspace dimension. Projecting the source activities onto these *Q* matrices and re-predicting target activities gave the performance when sequentially removing predictive dimensions.

Formally, we computed *X*_k_ = *XQ*, where:

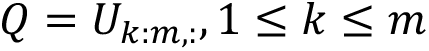

We then predicted performance from *X*_k_ to *Y* using ridge regression. Comparing to full-model performance revealed the effect of removing leading dimensions from communication subspaces.

### Canonical Correlation Analysis (CCA)

Canonical correlation analysis (CCA) was used to identify correlated activity patterns between the hippocampal CA1 and prefrontal cortex (PFC) populations. CCA finds linear combinations of the two populations that maximize correlation, known as canonical variates (Hotelling 1936; Steinmetz et al. 2019). In particular, we used CCA to continuously sample the overall communication subspace in a temporally resolved manner.

Spiking activity was binned in 50ms windows. CCA was applied to the binned spike count matrices of the CA1 and PFC populations to derive canonical variates that capture maximally correlated activity between the regions. 5-fold cross-validation was used to generate robust canonical variates (Steinmetz et al. 2019).

For examining individual canonical variates (Figures 2 and 6), the top 3 variates explaining the most correlated activity were analyzed. For behavioral prediction analyses (Figure 5), the top 10 canonical variates were used, below the typical optimal communication subspace dimension found through other analyses (Figure 4). Figure 4 predicted the 1^st^ CCA component using network patterns or speed.

For each canonical variate dimension, CCA produces a temporal activity pattern for CA1 (U) and PFC (V). The *U* and *V* temporal patterns were used to calculate aligned activity along the *U* = *V* unity line as (*U* + *V*)/*sqrt*(2), capturing reciprocated activity between areas. Orthogonal activity was calculated as (*U* − *V*)/*sqrt*(2) to measure unreciprocated activity.

### Network Pattern Distances

The Frobenius norm was computed on communication subspace matrix differences to quantify the distance and separation between them. The Frobenius norm measures the size of a matrix as the square root of the sum of their squared elements:

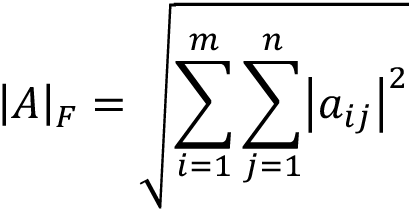

Where *A* is an *m* × *n* matrix and β_$7_ are its elements.

To examine distances between communication subspaces, the Frobenius norm was calculated between two *β* matrices *i* and *j* as:

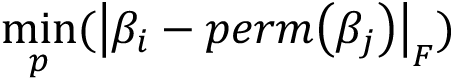

Where *perm*(*β*_j_) represents *β*_j_ with its columns randomly permuted. This permutation was repeated *p* = 100 times for each subspace pair to ensure the diversity of neuron pairings.

The norm was computed for all combinations of *β_j_* and *β_j_* from the set of subspaces for different high network pattern windows (**Figure 3 – Supplement 2**). The distribution of norms quantified the distances between subspaces.

### Behavioral Prediction with Linear Regression

Linear regression models were trained to predict rat behaviors from hippocampal-prefrontal communication space activity patterns derived with canonical correlation analysis (CCA). Models were constructed using TuringGLM.jl, a Julia package for Markov chain Monte Carlo sampled generalized linear models (GLMs) (Jose Storopoli and Rik Huijzer 2023). For each animal and behavior, 2000 MCMC samples were generated per GLM using the No U-Turn Sampler (NUTS) algorithm (Betancourt 2018).

50,000 random samples were used for training and the remaining held-out for testing, with separate GLMs per animal and behavior. Bernoulli regression was used for binary behavioral variables. Robust T-distributed regression was used for continuous variables to account for outliers.

The CCA source *U* and target *V* temporal activity patterns for the top 10 canonical variates were used as input features. These were projected onto an orthonormal aligned (reciprocated) and orthogonal aligned (unreciprocated) basis for the regression.

The models predicted velocity, acceleration, linearized position (0=center, 1=side wells), correct vs. error trials, outbound vs. inbound trajectories, integrated heading direction (IdPhi), and left vs. right trajectories.

For Figure 5 A-F, these models were computed using an entire session (day) of data. All animals with stable convergence of MCMC chains in all epochs were analyzed (N=6 animals out of 7). Epoch-wise models were created for Figure 5G, for which N=7 animals were used.

### Mixed Behavior and Local Field Potential Models

We created many different linear model specification formulas to test whether behavior, local field potential power, or some combination thereof better drove communication subspace activity. The goal was to understand whether one could attribute shared activity to one variable alone or the other, or if, instead, the activity requires both behavior and rhythmic network patterns.

In order to explore the information present in the parts versus the whole, we created data splits of animal, epoch, and each of three continuous behaviors (velocity, IdPhi, linear distance). In each split, we trained 2 models with either only behavior predicting the top shared activity component (*R* ∼ *B*) or theta local field power predicting the top shared activity component (*R* ∼ θ). And then we created 5 different models that incorporated some combination of these sources:

We assessed the model fit using metrics such as R-squared and AIC to ensure against overfitting for each model. We then compared the performance of these models to determine whether the inclusion of both behavior and theta power, as well as their interaction, provided a better fit to the data compared to models with only one predictor. This allowed us to assess the relative contributions of behavior and rhythmic network patterns in driving shared activity in the communication subspace and wholistic information beyond the parts indicated by higher *R*^2^ with lower AIC score on combined models.

### Theta Coherence Over Learning

To examine how theta coherence changed over learning, multiple coherence measures were calculated between CA1 and PFC as rats traversed the W-track across epochs. Coherence was quantified using both multi-taper coherence and weighted phase lag index (WPLI). WPLI provides increased robustness to noise confounds.

For both coherence measures, normalization was performed in two ways: 1) z-scoring across time within frequency bands per animal, and 2) min-max scaling per animal to a 0-1 range per frequency using scikit-learn applied to each animal’s full dataset. Data was bootstrapped such that each animal contributed equal samples to a given bootstrap. These two procedures correct for animals differing in measurement magnitude and for imbalances in number of temporal samples for animals.

To assess changes over learning, matched random subsets of times were sampled from each animal per epoch or spatial bin to ensure equal weighting across animals. Mean coherence and 95% confidence intervals were calculated from 1000 resampled subsets.

Spatial binning examined coherence differences across track locations. Analyses also evaluated coherence changes over learning epochs.

### Communication-Rhythm Coupling Over Learning

To quantify the relationship between communication space activity and theta power/coherence over learning epochs, linear regression models were trained. The input features were aligned, and orthogonal temporal activity patterns were found in the 1st canonical variate. The output being predicted was either theta power or coherence. Standard linear regression was used to model the relationship and obtain descriptive statistics. One model was fitted per epoch to capture relationship changes over learning.

Model performance was summarized by the coefficient of determination (*R*^2^). To obtain a population estimate, 10 subsets were randomly sampled and modeled per animal. Mean *R*^2^ and 95% confidence intervals were calculated across animals from these subsets to quantify how the linear relationship changed over epochs.

### Geometry of Communication Subspace

To investigate the relationship between neural activity and behavior, we constructed a pseudo-population representation by pooling data across multiple animals. Spiking activity from simultaneously recorded hippocampal and prefrontal neurons was binned according to behavioral variables, including trajectory direction (inbound/outbound), turn direction (left/right), trajectory (of in-out/left-right) and linear distance along the track. For each unique combination of these behavioral variables, we calculated the mean firing rate of each neuron across all animals. We then ordered by relative trajectory within trajectory type (in-out/left-right) and concatenated neurons across animals. This resulted in a super animal firing rate matrix, where each row represented a unique behavioral state, and each column represented a neuron. We then performed Canonical Correlation Analysis (CCA) on the super animal data, separating the activity into hippocampal and prefrontal components. This allowed us to identify patterns of co-activation between the two brain regions and to visualize the shared and unique contributions of each region to the overall neural representation of behavior. The resulting CCA components were further analyzed and visualized using dimensionality reduction techniques to explore the underlying structure of the neural-behavioral space.

### Software and Data Analysis

Data processing and communication subspace measurements were performed using MATLAB (version 9.14, The MathWorks, Inc., Natick, Massachusetts, United States). Tidy data structures were visualized using Python (version 3.9, Python Software Foundation, https://www.python.org/) with the following libraries: Matplotlib (version 3.6) (Hunter 2007), Seaborn (version 0.12) (Waskom 2021), and Pandas (version 1.5.2) (McKinney 2010). Markov Chain Monte Carlo (MCMC) methods were implemented in Julia (version 1.10) (Bezanson et al., 2017) using the TuringGLM.jl packages previously described.

## Data and Code Availability Statement

The data supporting the findings of this study are available in the Dandi Archive under the accession number DANDI:000978 Dandi Archive Link. The dataset includes all relevant recordings and metadata from the experiments described in this paper.

The code used for analysis and genera-on of results in this study is available on GitHub at hHps://github.com/jadhavlab/ca1pfc_commsub. Detailed instructions for data processing and analysis are provided within the repository.

## References

Abadchi, J. Karimi, Mojtaba Nazari-Ahangarkolaee, Sandra Gattas, Edgar Bermudez-Contreras, Artur Luczak, Bruce L. McNaughton, and Majid H. Mohajerani. 2020. “Spatiotemporal Patterns of Neocortical Activity around Hippocampal Sharp-Wave Ripples.” eLife. Retrieved November 2, 2023 (https://elifesciences.org/articles/51972/figures).

Barbosa, Joao, Rémi Proville, Chris C. Rodgers, Michael R. DeWeese, Srdjan Ostojic, and Yves Boubenec. 2023. “Early Selection of Task-Relevant Features through Population Gating.” Nature Communications 14(1):6837. doi: 10.1038/s41467-023-42519-5.

Battaglia, Francesco P., Karim Benchenane, Anton Sirota, Cyriel M. A. Pennartz, and Sidney I. Wiener. 2011. “The Hippocampus: Hub of Brain Network Communication for Memory.” Trends in Cognitive Sciences. doi: 10.1016/j.tics.2011.05.008.

Benchenane, Karim, Adrien Peyrache, Mehdi Khamassi, Patrick L. Tierney, Yves Gioanni, Francesco P. Battaglia, and Sidney I. Wiener. 2010. “Coherent Theta Oscillations and Reorganization of Spike Timing in the Hippocampal-Prefrontal Network upon Learning.” Neuron 66(6):921–36. doi: 10.1016/j.neuron.2010.05.013.

Benchenane, Karim, Paul H. Tiesinga, and Francesco P. Battaglia. 2011. “Oscillations in the Prefrontal Cortex: A Gateway to Memory and Attention.” Current Opinion in Neurobiology 21(3):475–85. doi: 10.1016/j.conb.2011.01.004.

Betancourt, Michael. 2018. “A Conceptual Introduction to Hamiltonian Monte Carlo.”

Bezaire, Marianne J., Ivan Raikov, Kelly Burk, Dhrumil Vyas, and Ivan Soltesz. 2016. “Interneuronal Mechanisms of Hippocampal Theta Oscillations in a Full-Scale Model of the Rodent CA1 Circuit” edited by F. K. Skinner. eLife 5:e18566. doi: 10.7554/eLife.18566.

Buzsáki, György. 2015. “Hippocampal Sharp Wave-Ripple: A Cognitive Biomarker for Episodic Memory and Planning.” Hippocampus 25(10):1073–1188. doi: 10.1002/hipo.22488.

Buzsáki, György, and Edvard I. Moser. 2013. “Memory, Navigation and Theta Rhythm in the Hippocampal-Entorhinal System.” Nature Neuroscience 16(2):130–38. doi: 10.1038/nn.3304.

Bygrave, Alexei M., Thomas Jahans-Price, Amy R. Wolff, Rolf Sprengel, Dimitri M. Kullmann, David M. Bannerman, and Dennis Kätzel. 2019. “Hippocampal–Prefrontal Coherence Mediates Working Memory and Selective Attention at Distinct Frequency Bands and Provides a Causal Link between Schizophrenia and Its Risk Gene GRIA1.” Translational Psychiatry 9(1):1–16. doi: 10.1038/s41398-019-0471-0.

Colgin, Laura Lee. 2016. “Rhythms of the Hippocampal Network.” Nature Reviews Neuroscience 17(4):239–49. doi: 10.1038/nrn.2016.21.

Colgin, Laura Lee, Tobias Denninger, Marianne Fyhn, Torkel Hafting, Tora Bonnevie, Ole Jensen, May-Britt Moser, and Edvard I. Moser. 2009. “Frequency of Gamma Oscillations Routes Flow of Information in the Hippocampus.” Nature 462(7271):353–57. doi: 10.1038/nature08573.

Dragoi, George, and György Buzsáki. 2006. “Temporal Encoding of Place Sequences by Hippocampal Cell Assemblies.” Neuron 50(1):145–57. doi: 10.1016/j.neuron.2006.02.023.

El-Gaby, Mohamady, Adam Loyd Harris, James C. R. Whittington, William Dorrell, Arya Bhomick, Mark E. Walton, Thomas Akam, and Tim E. J. Behrens. 2023. “A Cellular Basis for Mapping Behavioural Structure.”

Elsayed, Gamaleldin F., Antonio H. Lara, Matthew T. Kaufman, Mark M. Churchland, and John P. Cunningham. 2016. “Reorganization between Preparatory and Movement Population Responses in Motor Cortex.” Nature Communications 7(1):13239. doi: 10.1038/ncomms13239.

Fernández-Ruiz, Antonio, Azahara Oliva, Eliezyer Fermino de Oliveira, Florbela Rocha-Almeida, David Tingley, and György Buzsáki. 2019. “Long-Duration Hippocampal Sharp Wave Ripples Improve Memory.” Science (New York, N.Y.) 364(6445):1082–86. doi: 10.1126/science.aax0758.

Fernández-Ruiz, Antonio, Azahara Oliva, Gergő A. Nagy, Andrew P. Maurer, Antal Berényi, and György Buzsáki. 2017. “Entorhinal-CA3 Dual-Input Control of Spike Timing in the Hippocampus by Theta-Gamma Coupling.” Neuron 93(5):1213–1226.e5. doi: 10.1016/j.neuron.2017.02.017.

Floresco, S. B., J. K. Seamans, and A. G. Phillips. 1997. “Selective Roles for Hippocampal, Prefrontal Cortical, and Ventral Striatal Circuits in Radial-Arm Maze Tasks with or without a Delay.” The Journal of Neuroscience: The Official Journal of the Society for Neuroscience 17(5):1880–90.

Fries, Pascal. 2015. “Rhythms for Cognition: Communication through Coherence.” Neuron 88(1):220–35. doi: 10.1016/j.neuron.2015.09.034.

Gallego, Juan A., Matthew G. Perich, Lee E. Miller, and Sara A. Solla. 2017. “Neural Manifolds for the Control of Movement.” Neuron 94(5):978–84. doi: 10.1016/j.neuron.2017.05.025.

Haegens, Saskia, Yuriria Vázquez, Antonio Zainos, Manuel Alvarez, Ole Jensen, and Ranulfo Romo. 2014. “Thalamocortical Rhythms during a Vibrotactile Detection Task.” Proceedings of the National Academy of Sciences 111(17). doi: 10.1073/pnas.1405516111.

Hasz, Brendan M., and A. David Redish. 2020. “Spatial Encoding in Dorsomedial Prefrontal Cortex and Hippocampus Is Related during Deliberation.” Hippocampus 30(11):1194–1208. doi: 10.1002/hipo.23250.

Herbert, Elizabeth, and Srdjan Ostojic. 2022. “The Impact of Sparsity in Low-Rank Recurrent Neural Networks.” PLOS Computational Biology 18(8):e1010426. doi: 10.1371/journal.pcbi.1010426.

Hotelling, Harold. 1936. “Relations Between Two Sets of Variates.” Biometrika 28(3/4):321–77. doi: 10.2307/2333955.

Hunter, John D. 2007. “Matplotlib: A 2D Graphics Environment.” Computing in Science & Engineering 9(03):90–95. doi: 10.1109/MCSE.2007.55.

Hyman, James M., Bradley P. Wyble, Vikas Goyal, Christina A. Rossi, and Michael E. Hasselmo. 2003. “Stimulation in Hippocampal Region CA1 in Behaving Rats Yields Long-Term Potentiation When Delivered to the Peak of Theta and Long-Term Depression When Delivered to the Trough.” The Journal of Neuroscience: The Official Journal of the Society for Neuroscience 23(37):11725–31. doi: 10.1523/JNEUROSCI.23-37-11725.2003.

Hyman, James M., Eric A. Zilli, Amanda M. Paley, and Michael E. Hasselmo. 2005. “Medial Prefrontal Cortex Cells Show Dynamic Modulation with the Hippocampal Theta Rhythm Dependent on Behavior.” Hippocampus 15(6):739–49. doi: 10.1002/hipo.20106.

Iwata, Ryo, Hiroshi Kiyonari, and Takeshi Imai. 2017. “Mechanosensory-Based Phase Coding of Odor Identity in the Olfactory Bulb.” Neuron 96(5):1139–1152.e7. doi: 10.1016/j.neuron.2017.11.008.

Iyer, Ramakrishnan, Joshua H. Siegle, Gayathri Mahalingam, Shawn Olsen, and Stefan Mihalas. 2021. *Geometry of Inter-Areal Interactions in Mouse Visual Cortex*. *preprint*. Neuroscience. doi: 10.1101/2021.06.09.447638.

Jadhav, Shantanu P., Caleb Kemere, P. Walter German, and Loren M. Frank. 2012. “Awake Hippocampal Sharp-Wave Ripples Support Spatial Memory.” Science 336(6087):1454–58. doi: 10.1126/science.1217230.

Johnson, Niles G., David T. Guarrera, Homer F. Wolfe, DaGang Yang, and Morris Kalka. 2003. “The Riemannian Metric and Curvature Tensor on a Manifold.” 2(1).

Jones, Matthew W., and Matthew A. Wilson. 2005a. “Phase Precession of Medial Prefrontal Cortical Activity Relative to the Hippocampal Theta Rhythm.” Hippocampus 15(7):867–73. doi: 10.1002/hipo.20119.

Jones, Matthew W., and Matthew A. Wilson. 2005b. “Theta Rhythms Coordinate Hippocampal-Prefrontal Interactions in a Spatial Memory Task.” PLoS Biology 3(12):e402. doi: 10.1371/journal.pbio.0030402.

Joo, Hannah R., and Loren M. Frank. 2018. “The Hippocampal Sharp Wave-Ripple in Memory Retrieval for Immediate Use and Consolidation.” Nature Reviews. Neuroscience 19(12):744–57. doi: 10.1038/s41583-018-0077-1.

Jose Storopoli and Rik Huijzer. 2023. “TuringGLM.Jl.”

Joshi, Abhilasha, Eric Denovellis, Abhijith Mankili, Yagiz Meneksedag, Thomas Davidson, Anna K. Gillespie, Jennifer Ann Guidera, Demetris Roumis, and Loren M. Frank. 2022. “Dynamic Synchronization between Hippocampal Spatial Representations and the Stepping Rhythm.” 2022.02.23.481357.

Kaplan, Raphael, Mohit H. Adhikari, Rikkert Hindriks, Dante Mantini, Yusuke Murayama, Nikos K. Logothetis, and Gustavo Deco. 2016. “Hippocampal Sharp-Wave Ripples Influence Selective Activation of the Default Mode Network.” Current Biology 26(5):686–91. doi: 10.1016/j.cub.2016.01.017.

Kaufman, Matthew T., Mark M. Churchland, Stephen I. Ryu, and Krishna V. Shenoy. 2014. “Cortical Activity in the Null Space: Permitting Preparation without Movement.” Nature Neuroscience 17(3):440–48. doi: 10.1038/nn.3643.

Kim, Jaekyung, Abhilasha Joshi, Loren Frank, and Karunesh Ganguly. 2023. “Cortical–Hippocampal Coupling during Manifold Exploration in Motor Cortex.” Nature 613(7942):103–10. doi: 10.1038/s41586-022-05533-z.

Klausberger, Thomas, Peter J. Magill, László F. Márton, J. David B. Roberts, Philip M. Cobden, György Buzsáki, and Peter Somogyi. 2003. “Brain-State- and Cell-Type-Specific Firing of Hippocampal Interneurons in Vivo.” Nature 421(6925):844–48. doi: 10.1038/nature01374.

Kohn, Adam, Anna I. Jasper, João D. Semedo, Evren Gokcen, Christian K. Machens, and Byron M. Yu. 2020. “Principles of Corticocortical Communication: Proposed Schemes and Design Considerations.” Trends in Neurosciences 43(9):725–37. doi: 10.1016/j.tins.2020.07.001.

Kozachkov, Leo, Michaela Ennis, and Jean-Jacques Slotine. 2023. “RNNs of RNNs: Recursive Construction of Stable Assemblies of Recurrent Neural Networks.”

MacDowell, Camden J., Alexandra Libby, Caroline I. Jahn, Sina Tafazoli, and Timothy J. Buschman. 2023. “Multiplexed Subspaces Route Neural Activity Across Brain-Wide Networks.” 2023.02.08.527772.

Maharjan, Dennis M., Yu Y. Dai, Ethan H. Glantz, and Shantanu P. Jadhav. 2018. “Disruption of Dorsal Hippocampal – Prefrontal Interactions Using Chemogenetic Inactivation Impairs Spatial Learning.” Neurobiology of Learning and Memory 155:351–60. doi: 10.1016/j.nlm.2018.08.023.

McKenzie, Sam. 2018. “Inhibition Shapes the Organization of Hippocampal Representations.” Hippocampus 28(9):659–71. doi: 10.1002/hipo.22803.

McKinney, Wes. 2010. “Data Structures for Statistical Computing in Python.” Pp. 56–61 in. Austin, Texas.

Mitra, Partha, and Hemant Bokil. 2007. Observed Brain Dynamics. 1st edition. New York: Oxford University Press.

Moberly, Andrew H., Mary Schreck, Janardhan P. Bhattarai, Larry S. Zweifel, Wenqin Luo, and Minghong Ma. 2018. “Olfactory Inputs Modulate Respiration-Related Rhythmic Activity in the Prefrontal Cortex and Freezing Behavior.” Nature Communications 9(1):1528. doi: 10.1038/s41467-018-03988-1.

Mysin, Ivan. 2023. “Phase Relations of Interneuronal Activity Relative to Theta Rhythm.” Frontiers in Neural Circuits 17.

Nardin, Michele, Karola Kaefer, Federico Stella, and Jozsef Csicsvari. 2023. “Theta Oscillations as a Substrate for Medial Prefrontal-Hippocampal Assembly Interactions.” Cell Reports 42(9):113015. doi: 10.1016/j.celrep.2023.113015.

Nitzan, Noam, Rachel Swanson, Dietmar Schmitz, and György Buzsáki. 2022. “Brain-Wide Interactions during Hippocampal Sharp Wave Ripples.” Proceedings of the National Academy of Sciences 119(20):e2200931119. doi: 10.1073/pnas.2200931119.

Norman, Yitzhak, Omri Raccah, Su Liu, Josef Parvizi, and Rafael Malach. 2021. “Hippocampal Ripples and Their Coordinated Dialogue with the Default Mode Network during Recent and Remote Recollection.” Neuron 109(17):2767–2780.e5. doi: 10.1016/j.neuron.2021.06.020.

O’Neill, P. K., J. A. Gordon, and T. Sigurdsson. 2013. “Theta Oscillations in the Medial Prefrontal Cortex Are Modulated by Spatial Working Memory and Synchronize with the Hippocampus through Its Ventral Subregion.” Journal of Neuroscience 33(35):14211–24. doi: 10.1523/JNEUROSCI.2378-13.2013.

Papale, Andrew E., Jeffrey J. Stott, Nathaniel J. Powell, Paul S. Regier, and A. David Redish. 2012. “Interactions between Deliberation and Delay-Discounting in Rats.” Cognitive, Affective & Behavioral Neuroscience 12(3):513–26. doi: 10.3758/s13415-012-0097-7.

Redish, A. David. 2016. “Vicarious Trial and Error.” Nature Reviews Neuroscience 17(3):147–59. doi: 10.1038/nrn.2015.30.

Rezayat, Ehsan, Kelsey Clark, Mohammad-Reza A. Dehaqani, and Behrad Noudoost. 2022. “Dependence of Working Memory on Coordinated Activity Across Brain Areas.” Frontiers in Systems Neuroscience 15.

Santos-Pata, Diogo, and Paul F. M. J. Verschure. 2018. “Human Vicarious Trial and Error Is Predictive of Spatial Navigation Performance.” Frontiers in Behavioral Neuroscience 12:237. doi: 10.3389/fnbeh.2018.00237.

Schmidt, Brandy, Andrew Papale, A. David Redish, and Etan J. Markus. 2013. “Conflict between Place and Response Navigation Strategies: Effects on Vicarious Trial and Error (VTE) Behaviors.” Learning & Memory 20(3):130–38. doi: 10.1101/lm.028753.112.

Seger, Sarah E., Jennifer L. S. Kriegel, Brad C. Lega, and Arne D. Ekstrom. 2023. “Memory-Related Processing Is the Primary Driver of Human Hippocampal Theta Oscillations.” Neuron 111(19):3119–3130.e4. doi: 10.1016/j.neuron.2023.06.015.

Semedo, João D., Evren Gokcen, Christian K. Machens, Adam Kohn, and Byron M. Yu. 2020. “Statistical Methods for Dissecting Interactions between Brain Areas.” Current Opinion in Neurobiology 65:59–69. doi: 10.1016/j.conb.2020.09.009.

Semedo, João D., Amin Zandvakili, Christian K. Machens, Byron M. Yu, and Adam Kohn. 2019. “Cortical Areas Interact through a Communication Subspace.” Neuron 102(1):249–259.e4. doi: 10.1016/j.neuron.2019.01.026.

Shimbo, Akihiro, Ei-Ichi Izawa, and Shigeyoshi Fujisawa. 2021. “Scalable Representation of Time in the Hippocampus.” Science Advances 7(6):eabd7013. doi: 10.1126/sciadv.abd7013.

Shin, Justin D., and Shantanu P. Jadhav. 2016. “Multiple Modes of Hippocampal–Prefrontal Interactions in Memory-Guided Behavior.” Current Opinion in Neurobiology 40:161–69. doi: 10.1016/j.conb.2016.07.015.

Shin, Justin D., Wenbo Tang, and Shantanu P. Jadhav. 2019. “Dynamics of Awake Hippocampal-Prefrontal Replay for Spatial Learning and Memory-Guided Decision Making.” doi: 10.1016/j.neuron.2019.09.012.

Shin, Justin D., Wenbo Tang, and Shantanu P. Jadhav. 2023. “Protocol for Geometric Transformation of Cognitive Maps for Generalization across Hippocampal-Prefrontal Circuits.” STAR Protocols 4(3):102513. doi: 10.1016/j.xpro.2023.102513.

Siegle, Joshua H., and Matthew A. Wilson. 2014. “Enhancement of Encoding and Retrieval Functions through Theta Phase-Specific Manipulation of Hippocampus.” eLife 3:e03061.

Sigurdsson, Torfi, Kimberly L. Stark, Maria Karayiorgou, Joseph A. Gogos, and Joshua A. Gordon. 2010. “Impaired Hippocampal–Prefrontal Synchrony in a Genetic Mouse Model of Schizophrenia.” Nature 464(7289):763–67. doi: 10.1038/nature08855.

Somogyi, Peter, Linda Katona, Thomas Klausberger, Bálint Lasztóczi, and Tim J. Viney. 2014. “Temporal Redistribution of Inhibition over Neuronal Subcellular Domains Underlies State-Dependent Rhythmic Change of Excitability in the Hippocampus.” Philosophical Transactions of the Royal Society B: Biological Sciences 369(1635):20120518. doi: 10.1098/rstb.2012.0518.

Srinath, Ramanujan, Douglas A. Ruff, and Marlene R. Cohen. 2021. “Attention Improves Information Flow between Neuronal Populations without Changing the Communication Subspace.” Current Biology: CB 31(23):5299–5313.e4. doi: 10.1016/j.cub.2021.09.076.

Steinmetz, Nicholas A., Peter Zatka-Haas, Matteo Carandini, and Kenneth D. Harris. 2019. “Distributed Coding of Choice, Action and Engagement across the Mouse Brain.” Nature 576(7786):266–73. doi: 10.1038/s41586-019-1787-x.

Tang, Wenbo, Justin D. Shin, Loren M. Frank, and Shantanu P. Jadhav. 2017. “Hippocampal-Prefrontal Reactivation during Learning Is Stronger in Awake Compared with Sleep States.” The Journal of Neuroscience 37(49):11789–805. doi: 10.1523/JNEUROSCI.2291-17.2017.

Tang, Wenbo, Justin D. Shin, and Shantanu P. Jadhav. 2021. “Multiple Time-Scales of Decision-Making in the Hippocampus and Prefrontal Cortex.” eLife 10:e66227. doi: 10.7554/eLife.66227.

Tang, Wenbo, Justin D. Shin, and Shantanu P. Jadhav. 2023. “Geometric Transformation of Cognitive Maps for Generalization across Hippocampal-Prefrontal Circuits.” Cell Reports 42(3):112246. doi: 10.1016/j.celrep.2023.112246.

Tsodyks, Misha V., William E. Skaggs, Terrence J. Sejnowski, and Bruce L. McNaughton. 1996. “Population Dynamics and Theta Rhythm Phase Precession of Hippocampal Place Cell Firing: A Spiking Neuron Model.” Hippocampus 6(3):271–80. doi: 10.1002/(SICI)1098-1063(1996)6:3<271::AID-HIPO5>3.0.CO;2-Q.

Veuthey, T. L., K. Derosier, S. Kondapavulur, and K. Ganguly. 2020. “Single-Trial Cross-Area Neural Population Dynamics during Long-Term Skill Learning.” Nature Communications 11(1):4057. doi: 10.1038/s41467-020-17902-1.

Vinck, Martin, Francesco Paolo Battaglia, Thilo Womelsdorf, and Cyriel Pennartz. 2012. “Improved Measures of Phase-Coupling between Spikes and the Local Field Potential.” Journal of Computational Neuroscience 33(1):53–75. doi: 10.1007/s10827-011-0374-4.

Waskom, Michael. 2021. “Seaborn: Statistical Data Visualization.” Journal of Open Source Software 6(60):3021. doi: 10.21105/joss.03021.

Wilson, Matthew A., Carmen Varela, and Miguel Remondes. 2015. “Phase Organization of Network Computations.” Current Opinion in Neurobiology 31:250–53. doi: 10.1016/j.conb.2014.12.011.

Zhang, Haoxin, Ivan Skelin, Shiting Ma, Michelle Paff, Lilit Mnatsakanyan, Michael A. Yassa, Robert T. Knight, and Jack J. Lin. 2024. “Awake Ripples Enhance Emotional Memory Encoding in the Human Brain | Nature Communications -Https://Www.Nature.Com/Articles/S41467-023-44295-8.” *Nature Communications* 15(1):215. doi: 10.1038/s41467-023-44295-8.

Zhang, Yiyao, Liang Cao, Viktor Varga, Miao Jing, Mursel Karadas, Yulong Li, and György Buzsáki. 2021. “Cholinergic Suppression of Hippocampal Sharp-Wave Ripples Impairs Working Memory.” Proceedings of the National Academy of Sciences 118(15):e2016432118. doi: 10.1073/pnas.2016432118.

Zielinski, Mark C., Justin D. Shin, and Shantanu P. Jadhav. 2019. “Coherent Coding of Spatial Position Mediated by Theta Oscillations in the Hippocampus and Prefrontal Cortex.” Journal of Neuroscience 39(23):4550–65. doi: 10.1523/JNEUROSCI.0106-19.2019.

Zielinski, Mark C., Justin D. Shin, and Shantanu P. Jadhav. 2021. “Hippocampal Theta Sequences in REM Sleep during Spatial Learning.” 2021.04.15.439854.

Zielinski, Mark C., Wenbo Tang, and Shantanu P. Jadhav. 2020. “The Role of Replay and Theta Sequences in Mediating Hippocampal-Prefrontal Interactions for Memory and Cognition.” Hippocampus 30(1):60–72. doi: 10.1002/hipo.22821.

